# The ancestry of a genome or The amazing ubiquity of exponential distributions

**DOI:** 10.1101/2024.08.30.610585

**Authors:** Elizabeth Thompson

## Abstract

Motivated originally by the increasing number of examples where a lone member of a once thriving population or even species now survives, we investigate what genomes of that population the survivor may represent. More generally it is of interest to consider what genomes of our ancestry each of us may represent. We consider only diploid dioecious organisms, and consider primarily the ancestry of a haploid genome, for example the maternal autosomes of the focal individual. Our ancestors are many and, in an unbounded population, increase exponentially in number. Our genetic ancestors are few, bounded by the number of ancestral genome segments which increases linearly over past generations. First we show that the major loss of potential ancestral lineages is at 8-11 generations, and that thereafter the number of genetic ancestors increases approximately linearly, but does not approach the upper bound. Over many generations, there remain tightly linked but not contiguous segments that result from the same ancestral lineage. Second we analyze the process of these ”repeated” ancestral segments that continue to be formed in distant ancestry, even as others are lost by recombination. Thirdly, we consider the effect of a finite population, with one model of a geographically structured population. Ancestors are many, and soon fill the entire species range even with low migration rates. Genetic ancestors are not only few, but remain geographically local.

## 1 Introduction

There are an increasing number of highly endangered species where a single individual may be key to the genetic diversity of a future population to be reborn by captive breeding, or even cloning. These individuals may be the remnants of once thriving populations that existed tens of generations ago in Asia, North America, or Africa. Examples include the last wild-born Przewalski Horse captured as a foal in 1947, decades after the other members of her species had been living in captivity (Volf, 1961 et seq.; Ryder, 1988), or the last wild-born California condor captured in 1987, soon after the others of his species had been brought into captivity to ensure species survival (U.S. Fish & Wildlife Service, 1987). In both these cases, the species was numerically rescued if not restored, and wild populations reintroduced, but it is of interest to consider what genomes of their ancestral populations they may represent. More extreme examples include the two surviving female Northern White Rhinos (Spencer, 2020), and the Kew Garden cycad (Jones, 1993), the only known member, a male, of this dioecious diploid plant species.

In Section 2 we introduce the background and notation. We consider the autosomal genome of a diploid dioecious species, with *C* chromosomes and total genome length *L* Morgans. We focus on the ancestry of a single haploid genome, for example, the maternal genome of a current focal individual. Although the generation-to generation process of genetic recombination is straightforward, the process of ancestry is complex. Genetic ancestors are not only clustered within the tree of potential ancestral lineages, but ancestral lineages may occur more than once, even in an infinite population and over many generations. The purpose of this paper is to explore the processes that give rise to this clustering, and to explain why the number of genetic ancestors remains below the number of ancestral genome segments even in an unbounded population.

In Section 3 we discuss the clustering of surviving genetic lineages in the ancestry of a current individual, showing the clustering of such lineages, and the maximal loss of potential ancestral lineages at only 8 to 10 generations ago. The long-term branching pattern of lineages is also discussed. In Section 4 we look more directly at segments of ancestral genomes surviving in a current individual, and find how the number of genetic ancestors remains below the limiting bound of ancestral genome segments due to closely linked but not contiguous segments that are *repeats* of the the same remote ancestral genome. Because they are closely linked, with the surviving repeats ever more closely linked as time into the past progresses, these within-chromosome repeat segments remain over many generations.

In Section 5 we discuss the effect of finiteness of a population in decreasing the number of genetic ancestors. In a structured population (for illustration we use a stepping-stone model), we show that whereas an individual’s ancestors are many and everywhere, his genetic ancestors are not only few but also local. We conclude with a discussion Section 6.

## 2 Background and notation

### 2.1 Meiosis, crossovers, and recombination

In meiosis each chromosome of a diploid organism duplicates, and the maternal and paternal chromosomes align, forming the tetrad of four chromatids. At this stage, chiasmata form, resulting in the crossing over between homologous chromatids, leading to four potential offspring gametes. An example is shown in the upper part of Figure 1, with a resulting offspring chromosome consisting of three segments of the parent’s chromosome: two maternal segments with an intervening paternal segment.

**Figure 1:**
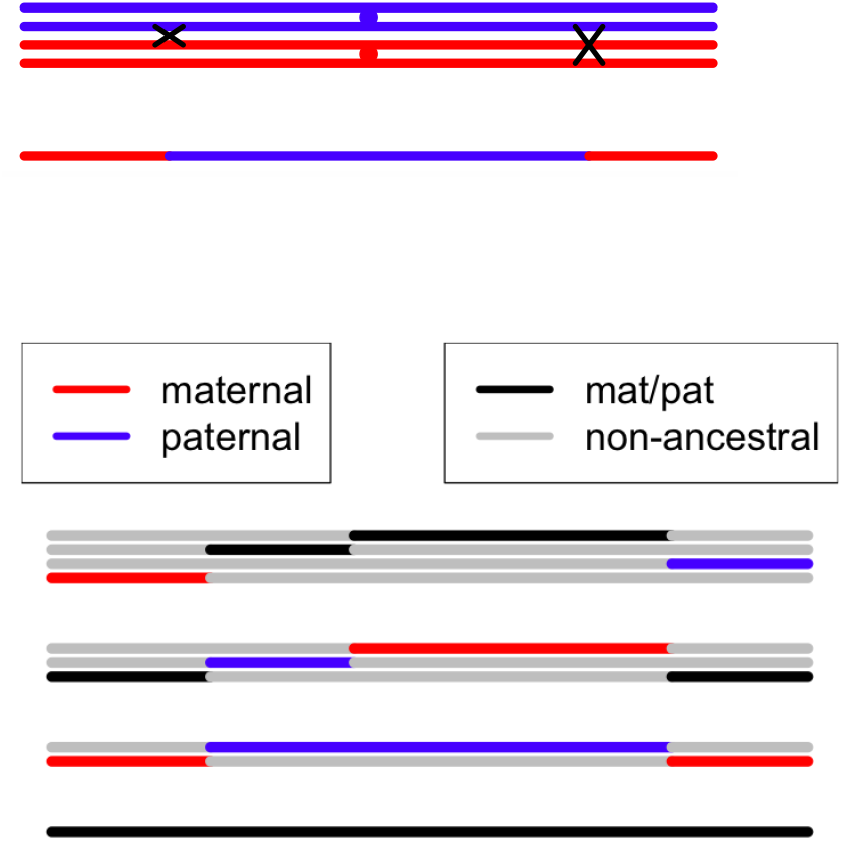
Above: crossovers occurring in a chromosome in meiosis. Below: the resulting processes of ancestry: see text for details.

Below is shown the process from the reverse time perspective. The final chromosome of an individual (shown in black) has ancestry in both the maternal and paternal chromosomes of the parent, with two maternal chromosome segments and one paternal.

Tracing back the earlier history of the segments shown in the lower part of Figure 1 there are two distinct processes that must be considered. First ancestral segments may be broken by crossover in the parental meiosis. For example the paternal segment above is shown splitting into a paternal (blue) and a maternal (red) segment of the parent’s parents. In this example it is shown that the chromosome with the two maternal segments was inherited intact from either the mother or the father of the initial parent (black). At the previous generation, the second relevant process is shown. Whereas the two central segments continue intact to either their maternal or paternal parents, the chromosome carrying two segments is broken by recombination (that is, an odd number of crossovers) in the *non-ancestral* intervening chromosome segment, so that the left-hand end derives from the maternal (red) chromosome of the parent, and the right-hand end from the paternal chromosome (blue). The four segments are now in four distinct ancestral genomes.

For the meiosis process we adopt the classical model of Haldane (1919), which assumes no chromatid interference, leading to a Poisson process of crossovers between maternal and paternal chromosomes in any offspring gamete. By definition, the rate of the Poisson process is 1 per Morgan. This leads to a very simple model for the splitting of ancestral segments. If a haploid genome (for example the maternal genome of a current individual) is traced backward into the ancestry of the individual, each of the *C* chromosomes will be broken by crossover events that occurred in the forwards transmission of DNA via meiosis. The total length of ancestral DNA will remain *L* Morgans, and the number of breakpoints occurring at each generation with be 𝒫*o*(*L*). Over *k* generations there will be 𝒫*o*(*kL*) breakpoints, and the total number of ancestral genome segments will be *C* + 𝒫*o*(*kL*), providing a simple upper bound on the number of distinct genetic ancestors of a current haploid genome.

The recombination process in non-ancestral material between ancestral segments on a single chromosome (Figure 1) is a little more complex, since recombination occurs only if there is an odd number of crossovers in the segment. The Poisson process of crossovers leads to the Haldane mapping function for recombination *ρ*(*d*) between two points at genetic distance *d* Morgans:

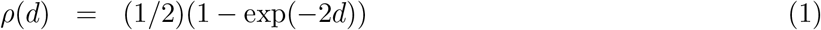

Successful meiosis requires at least one chiasma on the tetrad, so that any chromosome is passed intact to an offspring gamete with probability at most 1/2. That is, every autosome must have genetic length at least 0.5 Morgans, or 50 cM. The physically smallest chromosomes may have very small probability of more than one crossover in a single meiosis, rather than the value (1 − (3/2) exp(−0.5)) ≈ 0.09. given by the Poisson process. However, the assumption of the Haldane mapping function, or the potential for genetic interference will have little effect in our analysis, except possibly for the shortest chromosomes, since the vast majority of relevant crossovers will be occurring at different generations, and/or in different ancestors.

Note that everything in this paper relates only to genetic distance. The relationship between genetic and physical (base-pair) distance varies by species, chromosome location and other factors. As an overall average, 1 Morgan can be considered to be approximately 10^8^ base pairs. Exact results will depend on the number of chromosome pairs (*C*) and genetic length of the genome *L*, but results are qualitatively similar over a range of *C* and variation in lengths of individual chromosomes make up the total *L* (Thompson, 2023). For the purposes of numerical examples we assume an genome of 30 pairs of autosomes (*C* = 30) each of length 1 Morgan (*L* = 30): this approximates a wide variety of mammalian genomes.

### 2.2 The number and lengths of ancestral genome segments

For purposes of illustration, we simulate ancestry of 3000 chromosomes each of length 1 Morgan, simulated to a depth of 20 generations. Our pseudo-human genome consists of *C* = 30 such chromosomes, so that the haploid genome length is *L* = 30 Morgans. For the purposes of the current paper we divide the total set of chromosomes into just 100 sets of size 30, although the structure would allow for repeated random subdivisions of the 3000 chromosomes to allow better assessment of variance.

As noted above, for *C* chromosomes, and a genome of total length *L* Morgans, at *k* generations depth, there are *C* + 𝒫*o*(*kL*) segments ancestral to a single haploid genome: for example, the maternal autosomal genome of a current individual. Note this result does not depend on the lengths of individual chromosomes. Figure 2 shows the number of ancestral segments over generations 1 to 20. The 100 realizations are shown in grey; the mean in blue increases linearly in *k*, The standard deviation is 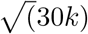, which increases slowly with *k*. The number of segments is, of course, an upper bound on the number of genomes that are genetic ancestors. The red curve shows the mean number of genetic ancestors at each generation: the reasons for the difference between red and blue lines will be discussed in Sections 2.3 and 4.2.

**Figure 2:**
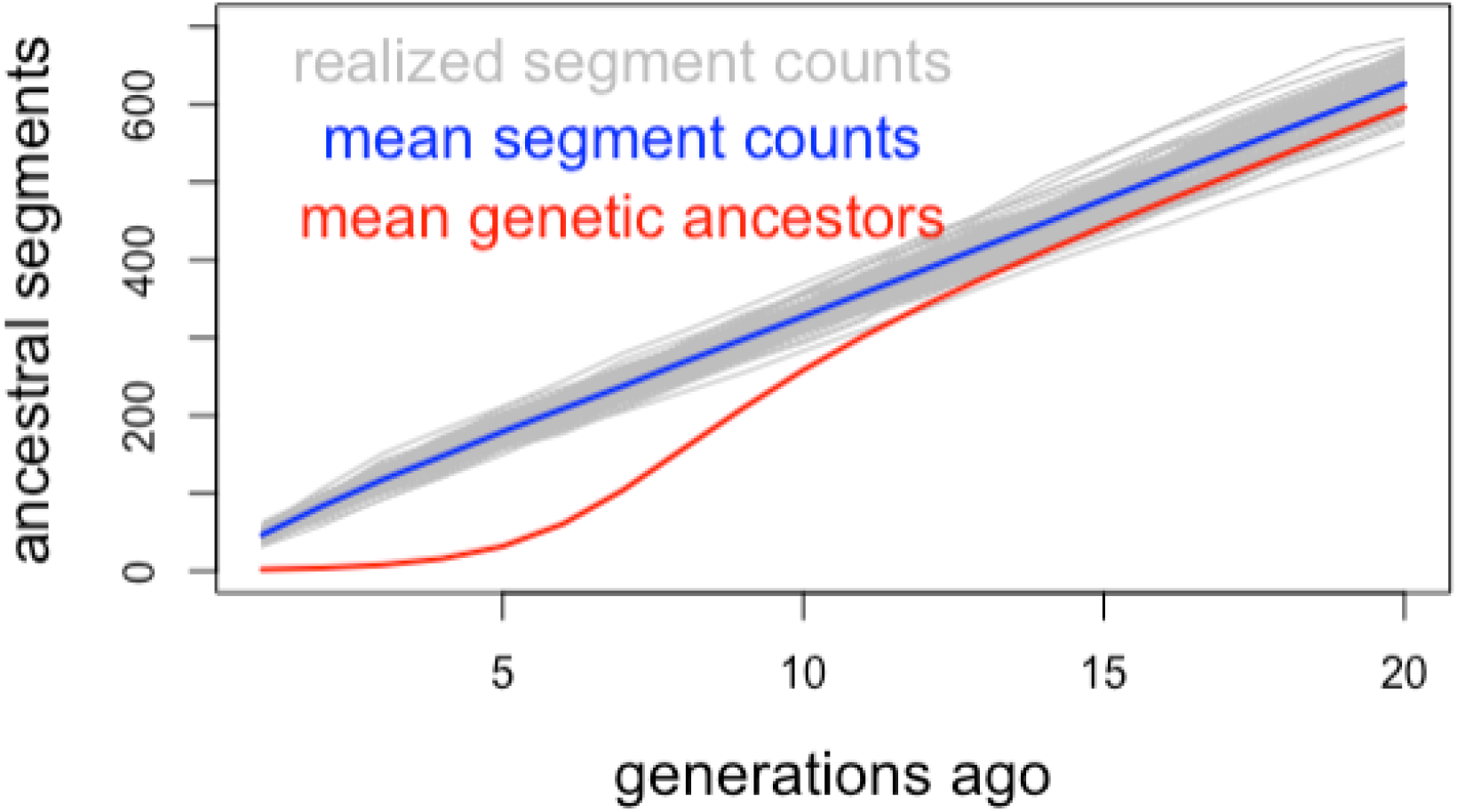
The number of ancestral genome segments over ancestral generations 1 to 20. The red curve shows the number of ancestral genomes represented by these segments, to which we return later. See text for details.

The distribution of lengths of the ancestral segments, in the absence of chromosome breaks, would be exponential with mean 1*/k*. Due to the chromosome breaks, the largest segments are lost, and the mean length is *L/*(*C* + *kL*), but there is little effect on the overall distribution. Figure 3 shows the segments arising in the simulated 100 genomes. For our case: at *k* = 20 theory gives a mean of 63,000 total segments in 3000 chromosomes of length 1 Morgan., with st.dev of 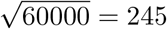 in the simulation we find 62,648. Unconstrained by chromosome breaks the expected length at *k* = 20 would be *k*^−1^ = 5 cM, but with the longest segments truncated the mean is *L/*(*C* + *kL*) or 4.76 cM. The simulation gives a mean segment length of 4.79 cM with a st.dev 4.76 cM; note, by chance there are slightly fewer than expected segments generated in this set of 3000 chromosomes.

**Figure 3:**
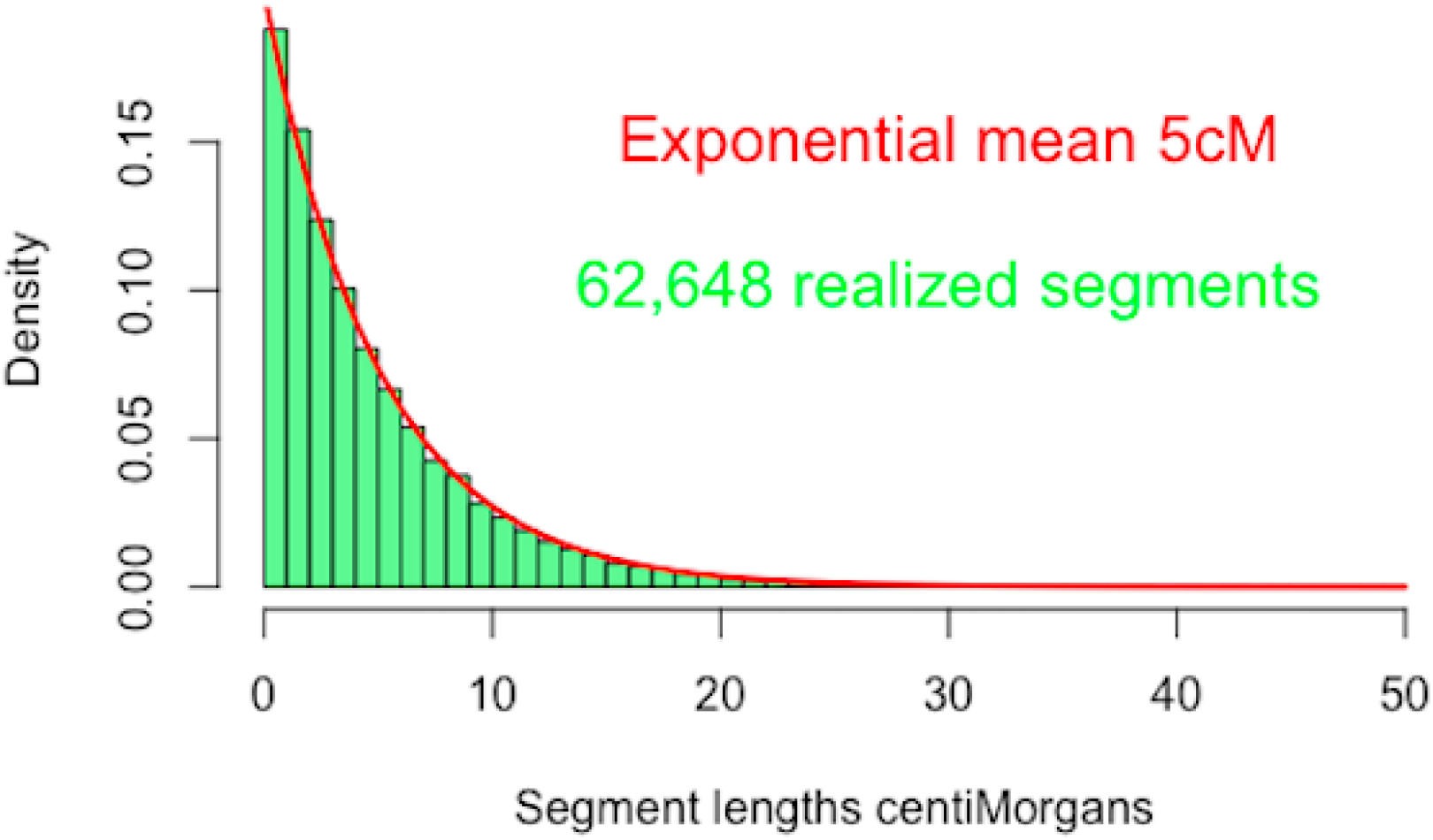
The lengths of 62,648 ancestral DNA segments at 20 generations depth.

We see how the exponential distrubution with many both very long and very short segments leads to counter-intuitive results. Disregarding the chromosome breaks, for the exponential with mean 5 cM. the probability of a segment length greater than 30 cM is exp(−6) = 0.0025, and 50cM in 4.5 × 10^−5^. At the other end of the scale, the probability of a segment less than 0.01 cM is (1 − exp(−0.05/5)) = 0.00200 almost equal to the frequency of segments of length greater than 30 cM. The exponential distribution of segment lengths results in the occurrence of more than one crossover within one ancestral segment in one meiosis, and hence the formation of ancestral chromosomes with two segments with the same ancestry (Figure 1), even at earlier generations.

The simulation of breakpoints on a chromosome is done in a continuum, but segments are recorded in units of 0.01 cM. In the 62,648 segments there are 8 recorded as length 0 (although the segment ancestry is still recorded), and 97 recorded as length 1. This gives a proportion 105/62648 = 0.00168 slightly less that than theoretical value. The longest segment is length is 46.3 cM, and the number greater than 30 cM is 120. The proportion 120/62648 = 0.00191 is very slightly lower than the continuous genome expectation, but clearly at *k* = 20 the effect of chromosomes on the segment length distribution is minimal.

### 2.3 The number of genetic ancestors

Figure 4 shows the results for lineages in more detail. The grey curves are now the realized number of distinct ancestral lineages in each 30-chromosome genome at generation *k*, and the red curve is the mean and is the same curve as shown in Figure 2. Note that the total number of segments is approximately Poisson and has mean (*C* + *kL*), and that marginally any segment has probability 2^*−k*^ of being a specified lineage. If lineages were independently assigned to segments, the number of segments of a specific lineage would be 𝒫*o*(2^*−k*^(*C* + *kL*)), and the probability a specified level-*k* ancestral genome does not occur in the current (*k* = 0) genome would be exp(−2^*−k*^(*C* + *kL*)). Thus the expected number of level-*k* lineages represented in the current genome would be

**Figure 4:**
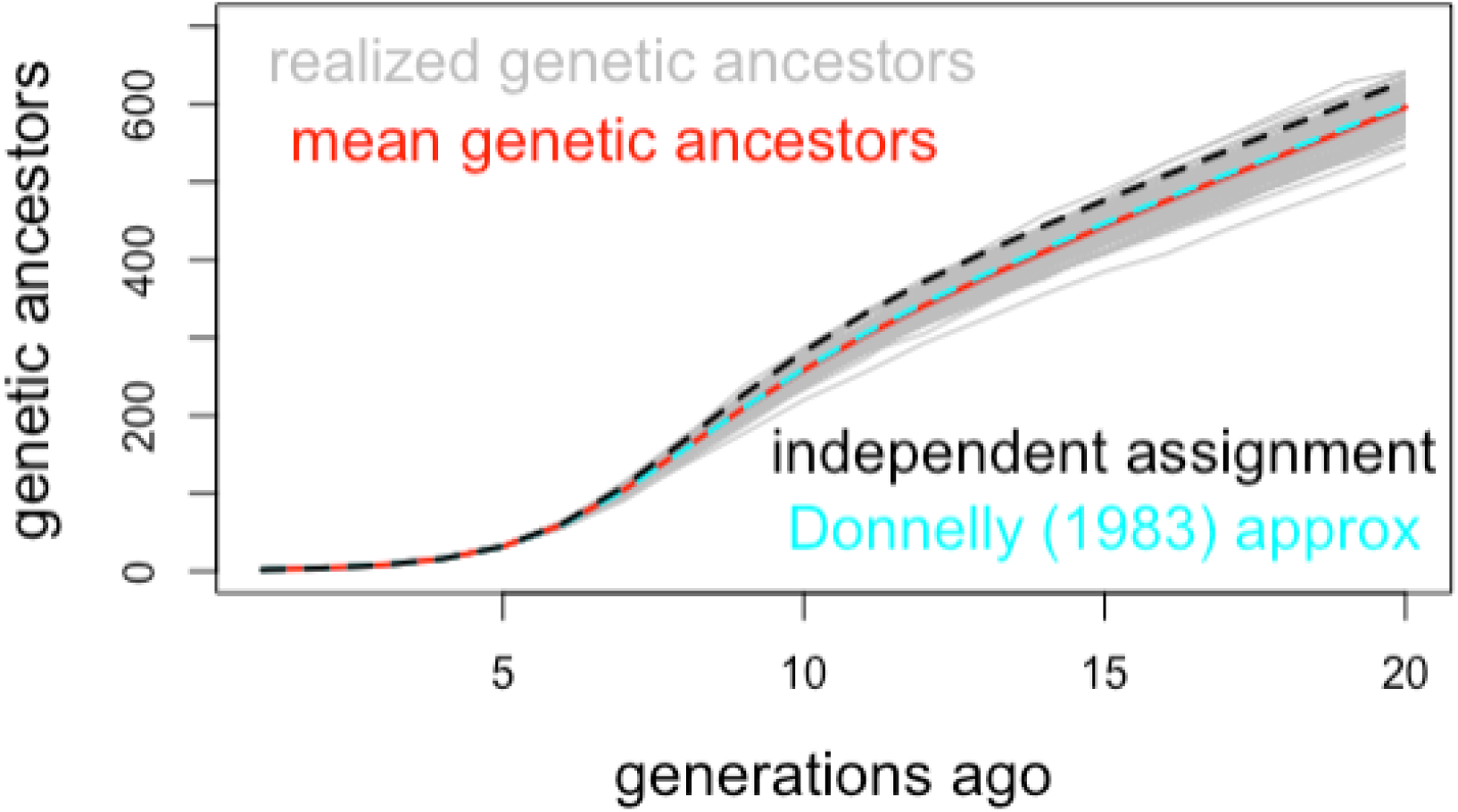
The number of distinct ancestral genomes (lineages) over ancestral generations 1 to 20. For details see text.

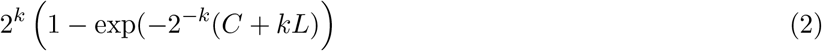

This is the approximation is the black-dashed line shown in Figure 4.

Note that for large *k* the value of formula (2) is approximately (*C* + *kL*), the expected total number of ancestral segments. This is, however, an overestimate of the number of ancestral genomes due to clustering of hits on any specific ancestral lineage across the genome. The clustering effect leading to repeated lineages remains even at larger *k* values. It was discussed by Donnelly (1983) and will be discussed further in Section 4.2. Donnelly (1983) gave a simpler approximation exp(−2^*−k*^*kL*) for the extinction of a lineage over *k* generations in a genome of length *L*. The expected number of lineages represented in a current genome would be 2^*k*^(1 − exp(−2^*−k*^*kL*). This approximation ignores both chromosome breaks and clustering of lineages, but gives a remarkably accurate approximation in many instances due to the compensating effect of these two approximations, in particular if *C* ≈ *L*. Donnelly’s curve is shown as the cyan dashed line in Figure 2 and is very close to the simulation mean. For the purposes of this paper we use the approximation exp(−*kL/*2^*k*^) developed by Donnelly (1983) for the extinction (non-survival) of an ancestral genome length *L* in a descendant at *k* generations remove.

## 3 Analysis of ancestral lineages

### 3.1 Notation: subtrees of potential ancestry

In this section we consider the *k*-generation haploid (maternal/paternal) ancestry of a single individual (Figure 5). There are 2^*k−*1^ diploid ancestors at generation *k*, and hence 2^*k*^ ancestor founder genomes (FGL) labelled 0,1,2,…, 2^*k*^ − 1. There is one meiosis from the mother (level 1) of the individual to the individual, 2 from her parents (level 2) to here, 4 from her grandparents (level 3) … There are 2^*k−*1^ meioses at level *k*, and a total of 2^*k*^ − 1 meioses in the ancestry. See Figure A.1 of Appendix A.1 for the case *k* = 3. All meioses switch independently 0 ↔ 1 at a rate 1 per Morgan. Note that the process here is across the chromosome, not backwards over generations.

**Figure 5:**
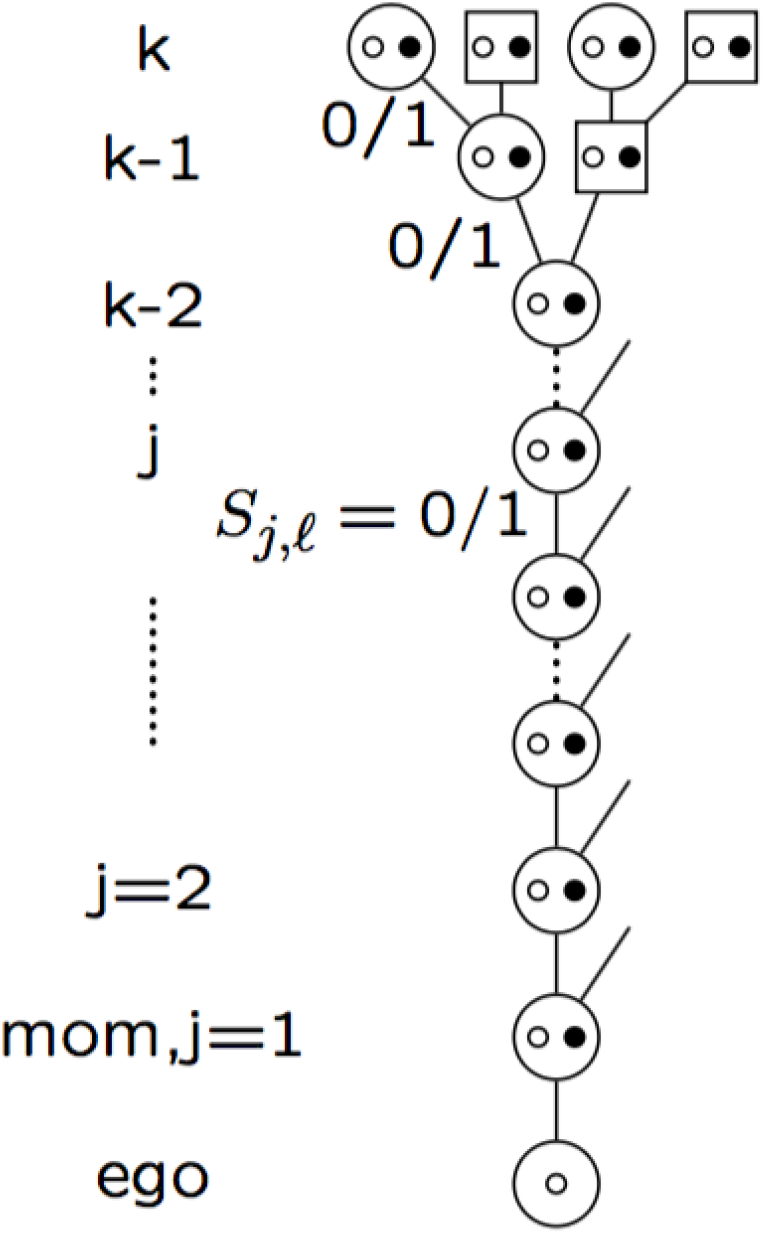
Schematic of k-generation maternal lineage ancestry of an individual

Donnelly (1983) frames the process of changing ancestry across a chromosome as a random walk on a hypercube. Each dimension of the cube corresponds to a meiosis in the ancestry, taking value either 0 or 1. The vertices define each 0/1 configuration of all meioses; there are 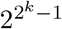vertices. In the case of direct ancestry, Donnelly (1983) considered just a single lineage of *k* meioses, but here we consider all the 2^*k*^ − 1 meioses in potential *k*-generation ancestries. We will label the meioses at level *j* as *S*_*j*,*l*_, for *j* = 1, …, *k*, ℓ = 1, …, 2^*j−*1^. Ancestral lineages may then be very simply notated as a sequence of *k* binary digits (0 and 1), and the lineages numbered by their decimal equivalents ranging from 0 = (00000 0) for the fully maternal lineage to 2^*k*^ − 1 = (1….1) for the fully paternal lineage, the lineages in 1-to-1 correspondence with the relevant ancestral FGL. This notation conveniently defines blocks of lineages rooted say at level *j* (*j* = 1, …, *k*), these lineages having the same first *j* binary digits, and comprising a block of 2^*k−j*^ integers in the decimal labelling.

At the start of chromosome, every meiosis is independently 0 or 1 with 50/50 probabilities. Each FGL then has probability 1/2^*k*^. A change in lineage occurs when one of the *k* meioses defining the lineage switches; each of the *k* lineages is the one to switch with probability 1*/k*. Note that the values of the other (2^*k*^ − 1 − *k*) meioses are irrelevant to a particular lineage. The total number of meioses patterns (vertices of the hypercube) is 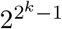 but these divide into 2^*k*^ lineages, with 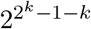 vertices corresponding to each FGL. If the meiosis at level *j* switches, the change will be to one of the 2^*k−j*^ FGL in the alternate subtree ancestral to that meiosis. In the absence of any other information, each node within that subtree will have equal probability 1/*k*2^*k−j*^. An example of the notation and setup for *k* = 3 is given in Appendix A.1.

Although the random walk on the full space of the vertices of the hypercube is Markov, the FGL process is not Markov, since previous states provide information on the values of some meioses. To proceed further we require the following key result:

#### Lemma

The probability any given meiosis is in changed state at the point at which the first of a disjoint set of *k* meioses changes state is 1/(*k* + 2).

Two derivations of this result are given in Appendix A.2. This result provides the probability 1/(*k* + 2) that any meiosis that was in the immediately preceding ancestral lineage, but not in the current ancestral lineage of *k* meioses is in changed state at the point on the chromosome at which the lineage changes again, and is sufficient to give two-step transition probabilities across a chromosome. The case *k* = 3 is worked through in Appendix A.3.

### 3.2 Transitions and subtree reversals

In general, knowledge of a transition between any two states that share ancestry up to level *j*_0_ provides information on the values of *j*_0_ + 2(*k* − *j*_0_) meioses at that point in the genome. The 2-step transitions for general *k* may be considered similarly to the case *k* = 3 (Appendix A.3). Again, for notational simplicity, we take the full maternal lineage and FGL 0 as the index state: *S*_*j*,1_ = 0, *j* = 1, …, *k*. To transition to FGL=0, the lineage of the previous state must differ by a single meiosis at level *j*_0_ say. Thus that state has lineage *S*_*j*,1_ = 0, *j* < *j*_0_, 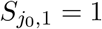, and *S*_*j*,𝓁_, for *j* = *j*_0_, …*k* determining the prior FGL. Only transition probabilities of FGL in the tree subtended at meiosis (*j*_0_, 1) are affected by the knowledge of the prior FGL,

At the next change of FGL there is equal probability 1*/k* of change in each *S*_*j*,1_. If the change is in 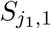 then for *j*_1_ ≠ *j*_0_ the probability is uniformly distributed over the 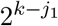 FGL in the relevant subtree rooted at level *j*_1_, since there is no prior information on the values of these meiosis switches. This is true whether *j*_1_ *< j*_0_, since then the subtree from 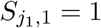 is disjoint from previous states. or *j*_0_ *< j*_1_, seen for example in the case of *k* = 3 for prior FGL 4 (*j*_0_ = 1), when *j*_1_ =2, resulting in equal probability for the FGL 6 and 7 (see Appendix A.3). If *j*_1_ = *j*_0_, 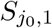 is the changed meiosis, and each meiosis in the lineage has changed state with probability 1/(*k* + 2) giving the distribution of the subtree as a collection of nested geometric probabilities to each successive subtree. The probability of reverting to the previous FGL is

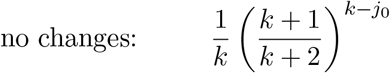

to the neighbor

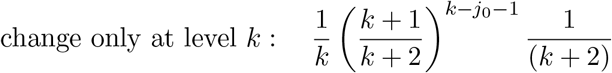

and so on.

Rather than considering transitions among individual states, it may be more relevant to consider subtrees rooted at a given level *j*_0_ say. A change at any level *j*_0_. changes the meioses defining the FGL lineage for all *j* > *j*_0_. If say 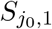 changes from 0 to 1, then subsequent changes in FGL at levels *j*_1_ > *j*_0_ do not affect any potential to revert to the previous FGL. The probability that 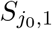 changes before any ancestral meioses is simply 1*/j*_0_, and return to the original subtree then occurs. The return will be to the previous state if the *k* − *j*_0_ meioses of the original state are unchanged at that point. For example consider the case *k* = 4 and the FGL sequence 2, 5, 6, 2:

| FGL | $S_{1,1}$ | $S_{2,1}$ | |
| --- | --- | --- | --- |
| $2 = (0, 0, 1, 0)$ | 0 | 0 | $S_{3,1} = 1 \quad S_{4,2} = 0$ |
| $5 = (0, 1, 0, 1)$ | 0 | 1 | $S_{3,2} = 0 \quad S_{4,3} = 1$ |
| $6 = (0, 1, 1, 0)$ | 0 | 1 | $S_{3,2} = 1 \quad S_{4,4} = 0$ |
| $2 = (0, 0, 1, 0)$ | 0 | 0 | $S_{3,1} = 1 \quad S_{4,2} = 0$ |

In this case *j*_0_ = 2, and the change from 0 to 1, moves the FGL from the set {0, 1, 2, 3} to the set {4, 5, 6, 7}, the alternate sets rooted at *S*_2,1_. The next change is within the latter subset as *S*_3,2_ changes from 0 to 1: note that different meioses are then relevant at level 4. These changes, and any others within the subtree, are irrelevant to any return to the FGL=2. The probability that *S*_2,1_ changes before *S*_1,1_ is 1*/j*_0_ = 1/2, In this case, the FGL will return to the set {0, 1, 2, 3}, and to the original FGL=2 provided both *S*_3,1_ and *S*_4,2_ are then unchanged. The sequence of FGL lineages across a chromosome is thus complex. And alternate view of returns to lineages across the chromosome will be presented in Section 4.2.

### 3.3 Clustering of surviving lineages

In considering lineages surviving in a current individual from say *k* generations ago, it is easier to consider the generations *j* (*j* ≤ *k*) increasing backwards in time, rather than the process across the chromosome. At *k* ≥ 15 for a genome length *L* approximately *kL* of the 2^*k*^ lineages survive but these are far from a random subset of the 2^*k*^ lineages: lineage loss shows strong patters of dependency. When, proceeding back in time, a genome is lost, say at generation *j*, then so are all 2^*k−j*^ lineages at generation *k* that are ancestral to it. Specifically, we illustrate the case *k* = 20, in the simulation of 100 genomes of *C* = 30 chromosomes each of length 1 Morgan (Section 2.2). At generation 5, a single lineage is lost in 16 of the 100 genome realizations, but each of these represents 2^15^ = 32, 768 ancestors at generation 20.

Figure 6 shows (on a log base-2 scale) the lineage losses in the set of 100 simulated 30-chromosome genomes. Grey lines (largely superimposed) give the 100 outcomes, and the blue line is the mean. Losses are accounted by the number of potential generation-20 ancestors eliminated by the loss. We see that the maximum loss occurs at generation 8 or 9, when larger numbers of new losses occur, each eliminating a block of several thousand generation-20 lineages. (Note that the log-base-2 of the means is distorted in the early generations:: log_2_(Mean(x)) ≠ Mean(log_2_(x)).)

**Figure 6:**
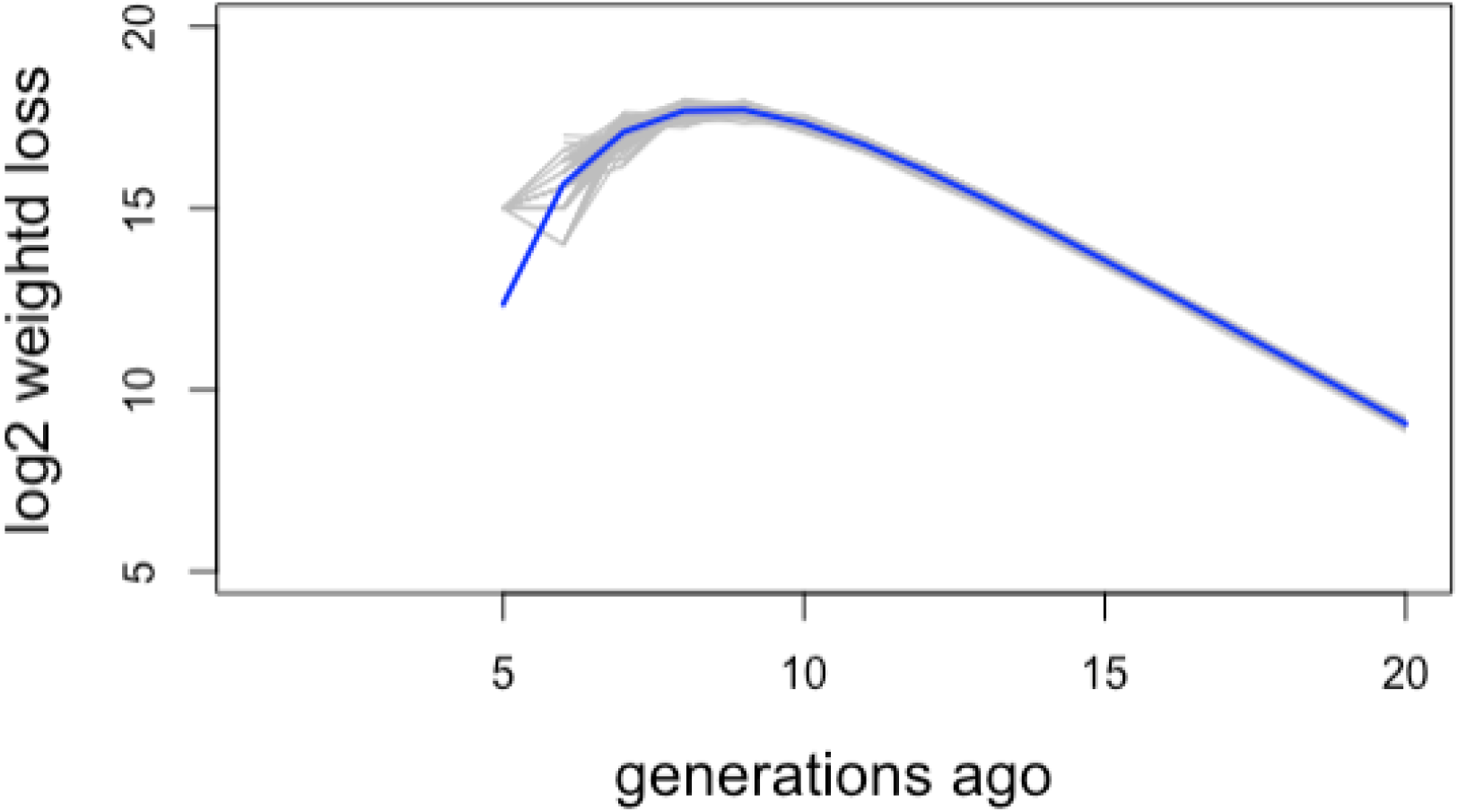
Ancestral lineage loss. See text for details.

We see that the first lineages are lost at generation 5, while by generation 16 or so the number of newly lost lineages is close to the number surviving. That is the surviving ancestral segment will usually come from one parent or the other. In fact the number when both survive is non-negligible. At every generation there will be a P*o*(*L*) number of breakpoints generated in the ancestry of the current genome, splitting an ancestral segment into the two lineages of the parent individual. That is, however large is *k*, there will be an average *L* = 30 ancestors at generation *k* both of whose genomes survive. This leads to strong positive correlation in survival of two genomes within a single ancestor as adjacent segments in the descendant (Thompson, 2023).

The cumulative distribution of the indices of the *k*^*th*^-generation ancestors with genome in the current index genome gives an alternative representation of the clustering of lineage loss. Figure 7 shows, for a single realized genome the empirical cumulative distribution (*ecdf*) of the indices of the 566 surviving genomes, among the 2^20^ = 1, 024, 576 potential ancestral lineages. Flat segments of the *ecdf* indicate blocks of lineages that are lost due to a single early loss. There is a large flat segment at approximately 3.6 to almost 4 × 10^5^, indicating an early loss: indeed the lineage 11 = (0, 1, 0, 1, 1) is lost at generation 5 in this realization: 11 × 2^15^ = 360, 448, 12 × 2^15^ − 1 = 393, 215. Smaller flat segments indicating later losses at around indices 6 to 8 × 10^5^. Conversely, steep segments of the curve shown where a cluster of ancestrally close lineages survive– two genomes within an individual, in co-parents, or in co-grandparents.

**Figure 7:**
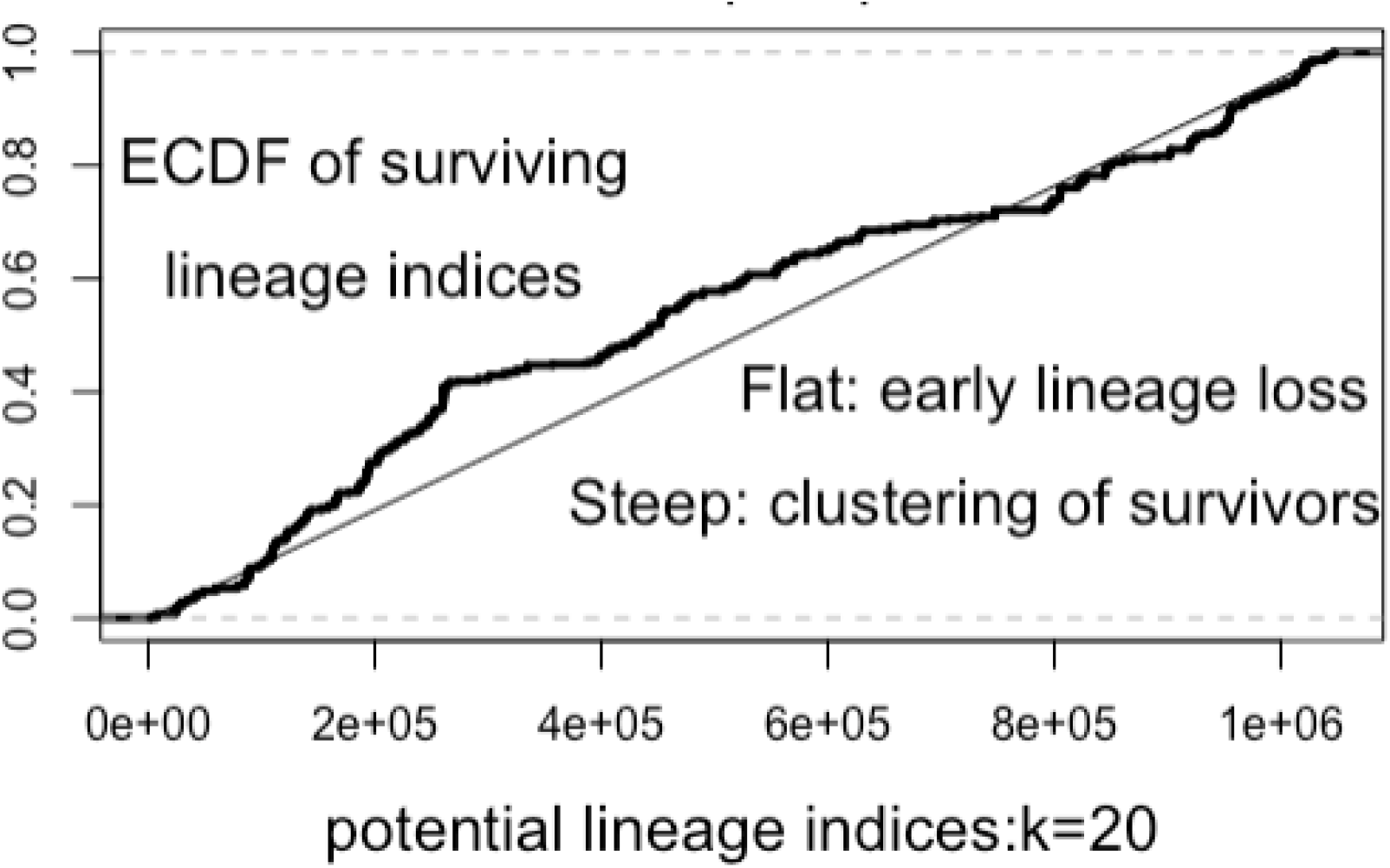
The ECDF of the 566 ancestral lineage labels with DNA surviving over 20 generations in one realization of a genome of 30 chromosomes length 1M.

### 3.4 Surviving lineage structure at greater time depths

For *k* ≥ 20, there are approximately *kL* genetic ancestors, and segments are marginally approximately exponential with mean 1*/k*. For the purposes of the current section we will ignore the truncation of segments by chromosome ends, which is increasingly irrelevant at *k* becomes large. We will also not consider separately genomes that carry more than one ancestral segment. We will find in Section 4.2 that for *k* ≥ 20 these will be repeated tightly linked segments on single chromosomes, so that for the current discussion on their continuing ancestry they will behave very similarly to genetic ancestors carrying a single ancestral segment. A genetic ancestor with ancestral genome length *ℓ* will have a single genetic-ancestor parent with probability exp(−*ℓ*) and both male and female parents will be genetic ancestors with probability *p*(*ℓ*) = (1 −exp(−*ℓ*)) The number of generations until this branching into two lineages occurs is Geometric with success probability *p*(*ℓ*). Over a subsequent *t* generations, the number of splits partitioning the segment is Poisson 𝒫*o*(*tℓ*), so the lineage will have 1 + 𝒫*o*(*tℓ*) ancestors surviving in generation 0 at generation *t* + *k*.

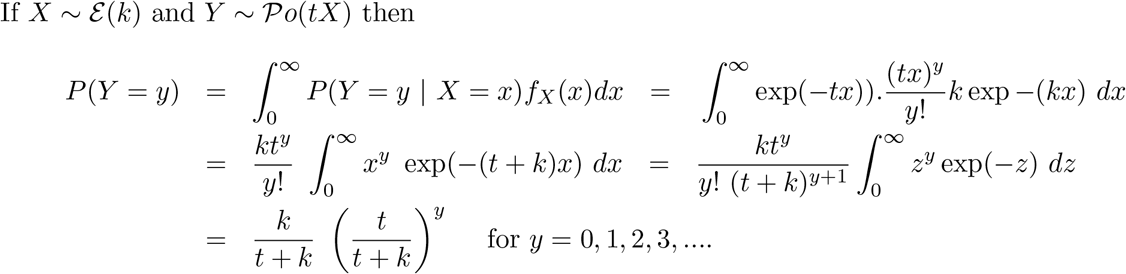

Thus overall the number of additional surviving lineages that a generation *k* lineage gives rise to at generation (*k* + *t*) has a geometric distribution, and the expectation increases linearly in *t*.

This overall geometric distribution is however slightly misleading as to the clusters of surviving lineages at greater time depths, since clearly the longer ancestral segments are the ones more likely to be broken, giving rise to two surviving parental lineages. Due to the exponential distribution of segment lengths (Figure 3) long segments remain even at generation *k* = 20 when the mean length is under 5 cM. Figure 8 shows the maximum length of an ancestral segment at *k* = 20 in each of the 3000 simulated chromosomes (left) and in the 100 genomes (right) formed from sets of 30 chromosomes. We see that each chromosome typically contains a segment at least 3 times the mean, while each genome typically has a segment of 30 cM or more. These longer segments offer opportunities for substantial branching of ancestry even at greater time depths. For example, a segment of 30cM length has probability over 25% of an immediate split to both the paternal and maternal genomes of its parents, and a probability 1% of more than 4 genetic ancestors within 5 generations.

**Figure 8:**
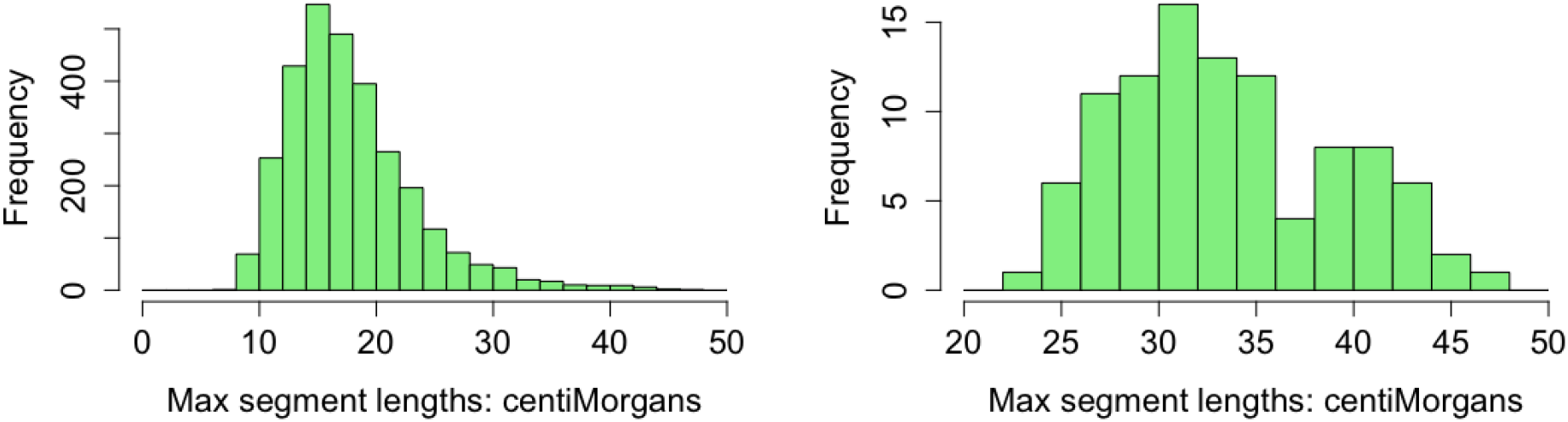
The maximum lengths of ancestral segments at generation *k* = 20. Left: Maximum in each of 3000 chromosomes. Right: Maximum in each of 100 genomes.

## 4 Clustering and repeated ancestral segments across the genome

### 4.1 Clustering of lineage indices of ancestral segments across the genome

We now consider, as did Donnelly (1983), the clustering of “hits” on a surviving ancestral lineage across the genome. Our problem is more complex than the framework of that paper. There the focus was on the lineage from a specific ancestor. Thus only the process of the *k* meioses to that ancestor and the clustering of hits on the specific lineage were considered. In contrast, we consider all the lineages to ancestors who contribute to the final genome.

As an example, Figure 9 shows, for the same example as shown in Figure 7 the lineages indices of the 607 ancestral segments at 20-generation-depth ancestry across the genome. (Note: 566 of the 607 lineages are distinct: cf Figure 7.) The chromosomes each have the same genetic length (1 Morgan), but different numbers of ancestral segments ranging from 12 (Chromosome-1) to 26 (Chromosome-11). We see that there are some early crossovers in ancestral segments leading to dispersed lineage indices on that chromosome. However, other chromosomes have indices within one half, or even one quarter of the range, indicating no splits in the first one or two generations. Generally successive indices are close in value, indicating splitting at deeper generations.

**Figure 9:**
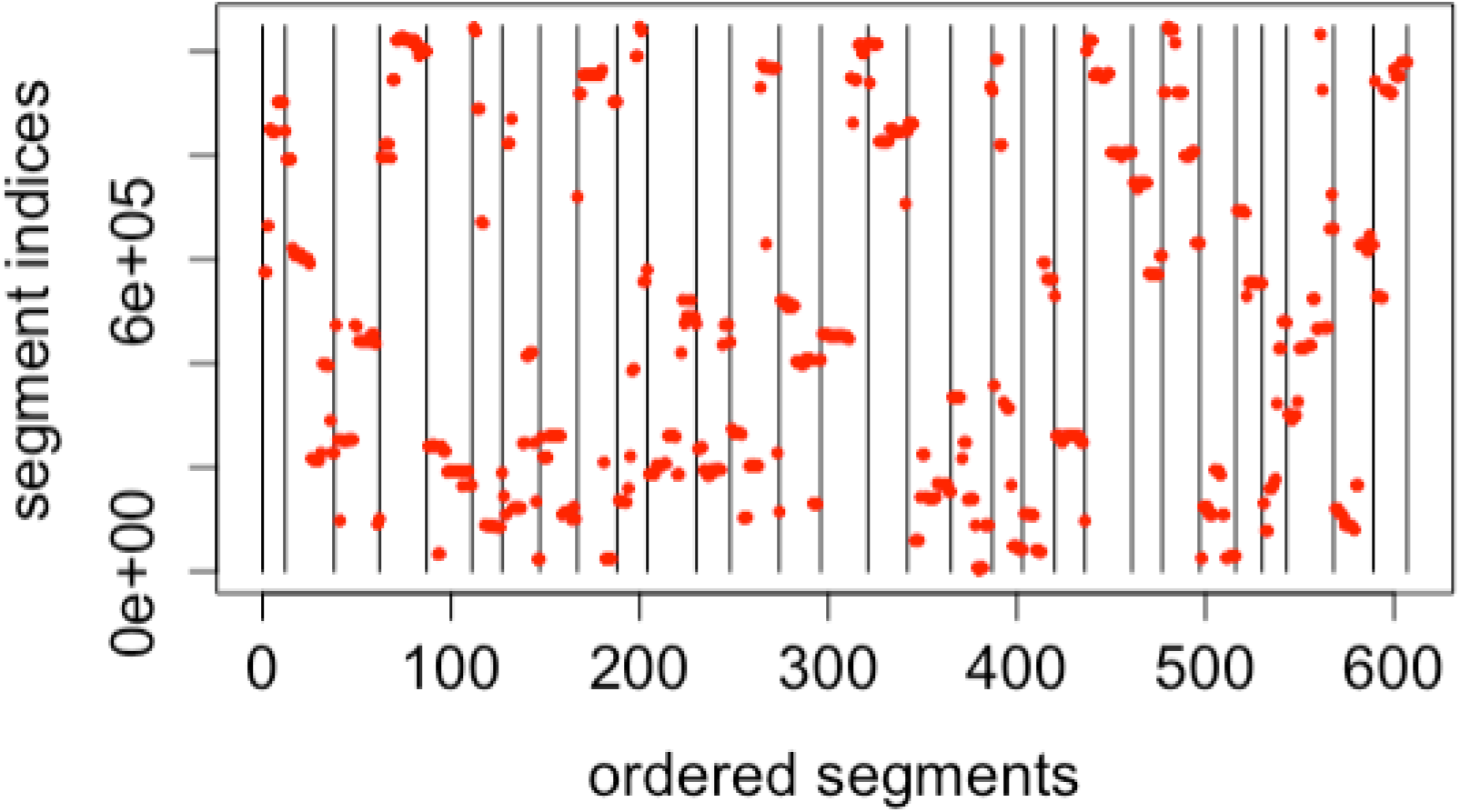
Example of the clustering of lineage indices of surviving ancestral segments at 20 generations depth within each chromosome, across the genome.

Note however then due to the binary indexing of the lineages, a split at generation 1 generates a change of index between the lower and upper halves of the graph, whereas, for example, a split at generation 5 is a change of only 1/2^5^ or about 3% of the vertical range. *A priori* an equal number of splits of ancestral material is expected at each generation, and in each chromosome. *A posteriori* any given split has equal chance or having occurred at any generation. This is shown in Figure 10 where the changes in lineages indices across each chromosome are shown on a log_2_ scale. Specifically, what is plotted on the vertical axis is

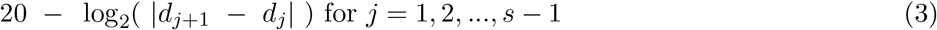

where *s* is the number of ancestral segments in the chromosome, and *d*_*j*_ is the decimal index of ancestral segment *j* in the chromosome. This represents the generation depths at which the ancestral segment is broken and the lineages diverge, and we see the approximate uniformity of these depths. Note that the generation 20 splits align exactly at height 20: the change in index will be exactly 1, and 20−log_2_(1) = 20. For other generations this is not the case. For example at generation 10, the first binary difference is in the 10th binary digit, but thereafter the next 10 binary digits in one ancestral lineage will be independent of those in the other, giving a range of values between 10 and 11 to the measure (3). Note also that several chromosomes have no splits in the first generations. This is unsurprising: at any generation the probability of no split of a 1 Morgan length is exp(−1) = 0.3679.

**Figure 10:**
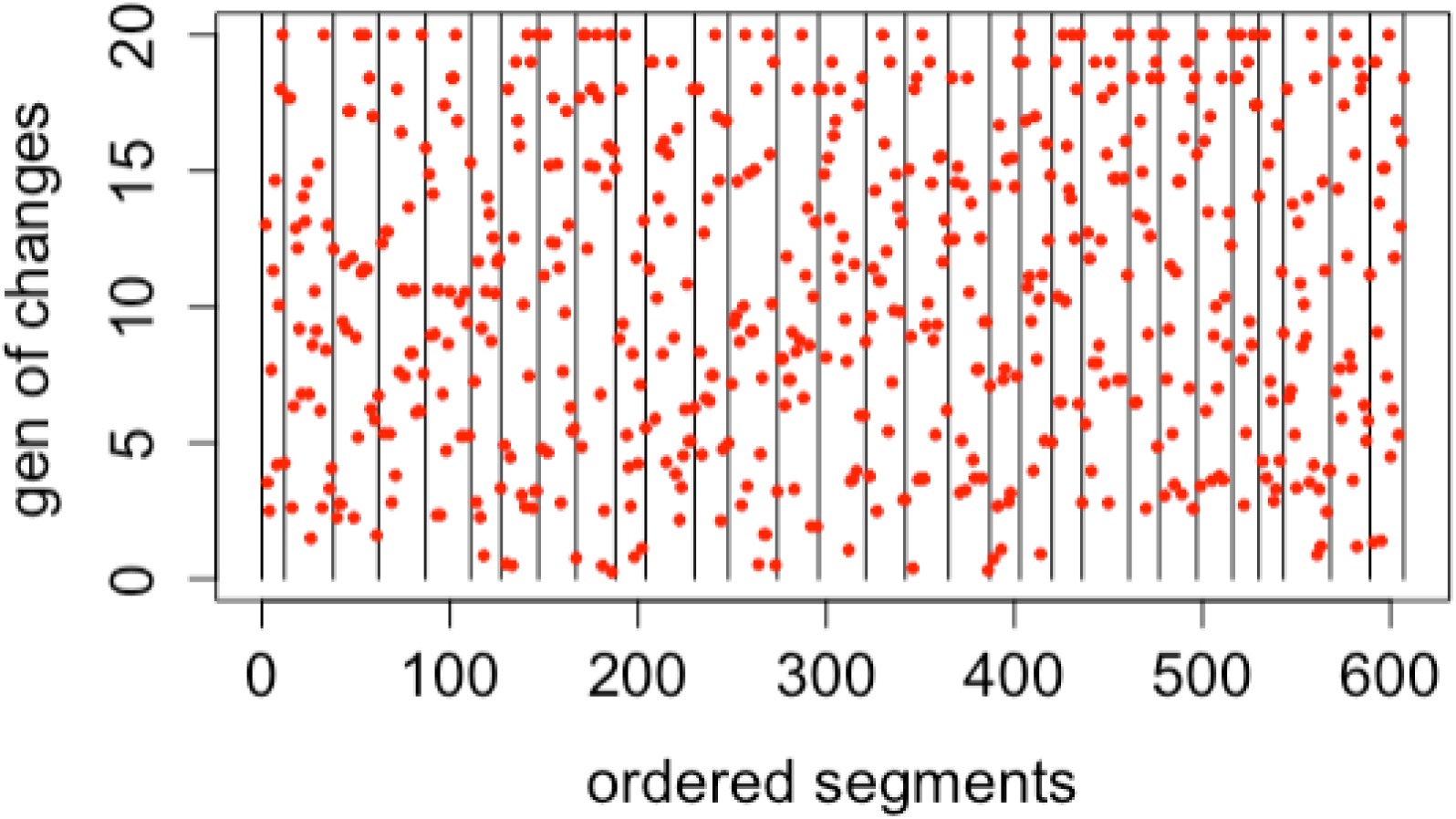
Differences in successive ancestral lineage indices on a log_2_ scale for ancestral segments at generation 20 across each chromosome, representing the generation depth of each transition.

### 4.2 Segments of repeated lineage indices between and across chromosomes

Figure 4 (see Section 2.3) shows that the number of distinct surviving lineages remains below the number of ancestral segments even at deeper generations. That is, some lineages are repeated. Within a chromosome, a repeat is formed when there is more than one crossover within an ancestral segment in a single meiosis.

This was seen in the lower part of Figure 1 repeated here as Figure 11. The two end segments of the maternal chromosome in the parental generation are separated by non-ancestral genetic material. These two maternal segments will remain on a single chromosome, and share ancestry, until a recombination in the non-ancestral segment separates them; in this small example this occurs at the top-most generation shown.

**Figure 11:**
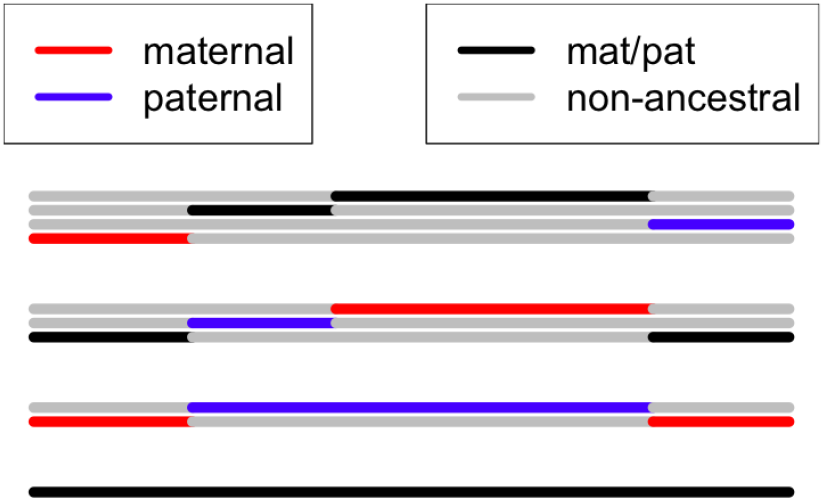
The creation and subsequent loss of a repeated segment with identical ancestry.

The central segment located on the paternal chromosome in the parental generation will diverge to an independent ancestry, possibly breaking into segments (as shown in the grandparents) and/or creating its own repeats within it. Because of the potential for the exponential distribution to produce both many short and some long crossover intervals, repeats can be generated even at deeper generations, and earlier ones can remain without being broken by recombination.

We show first the results of the example simulation of 100 genomes, each of 30 1-Morgan chromosomes. Figure 12 shows the total counts of segments than are lineage repeats across the 100 genome realizations.. There are a total of 62648 segments in all. Each segment that is a repeat of a lineage contributes once only to the count. For example, if within a genome (set of 30 chromosomes) a lineage occurs on 5 different chromosomes, including 3 times on 1 chromosome and 2 times on another, then the lineage contributes 5 − 1 = 4 to the cross-chromosome count, and (3 − 1) + (2 − 1) = 3 to the within chromosome repeat count. Up to 5 or 6 generations depth, there are many repeats of lineages both within and across chromosomes, but the cross-chromosome repeats decline rapidly as generation depth increases. By depth *k* = 20 there are very few cross-chromosome repeats, as ancestral paths diverge under the independent inheritance in each chromosome. In fact, there are just 17 in the total 100 genome simulations. By contrast, the number of within-chromosome repeats remains roughly constant, although declining very slightly. A total of 3036 remain at generation 20, or just over one per chromosome on average. The maximum was 3271 at generation 5. These counts confirm the discussion of Donnelly (1983) that the effect of chromosome breaks (*C* = 30) effectively cancels the repeats due to clustering in a genome length *L* = 30 Morgans.

**Figure 12:**
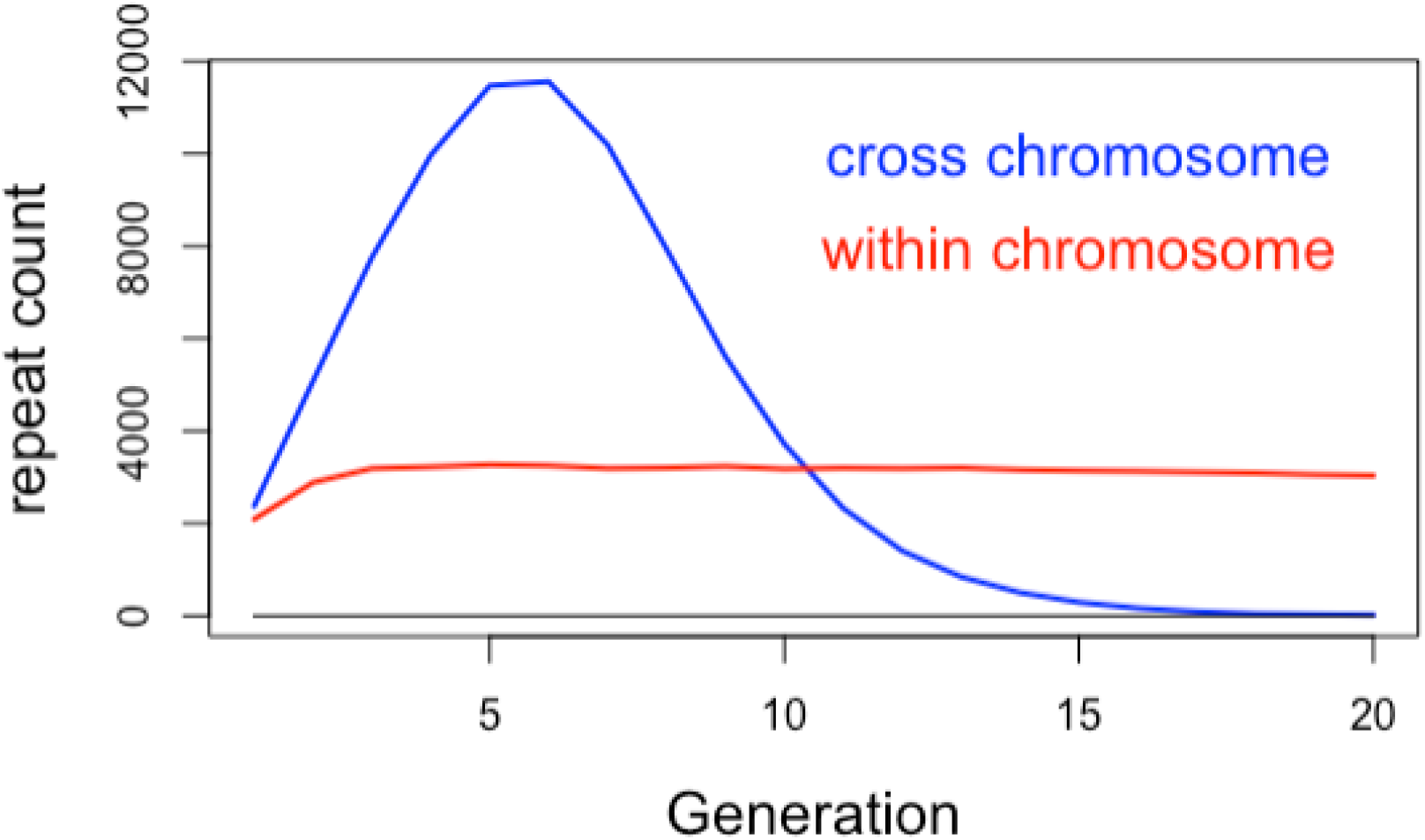
Total repeated lineage segments within genomes, across and within chromosomes over 20 generations.

We use the term *spacings* for the segments of non-ancestral DNA between within-chromosome *repeat* segments. Figure 13 shows the genetic distance lengths of spacings. Again each repeat contributes once only. For example, if a lineage occurs in 3 segments on a chromosome, the distances are the length of the spacing between the first and second occurrence, and then between the second and third. At 5 and 10 generations depth, there is little evidence of clustering of hits within a chromosome, but by depth *k* = 15 most spacing lengths are small, and more so by *k* = 20. The proportion of lengths that are less than 5 cM are 20.4%, 35.8%, 49.4%, and 60.3% over depths *k* = 5, 10, 15 and 20. The mean distances are 23.3 cM, 12.8 cM, 8.2 cM and 5.8 cM, respectively. As *k* increases, fewer repeats are formed as ancestral segments are smaller, but the decreasing lengths of spacings mean that repeats remain for more generations into the past.

**Figure 13:**
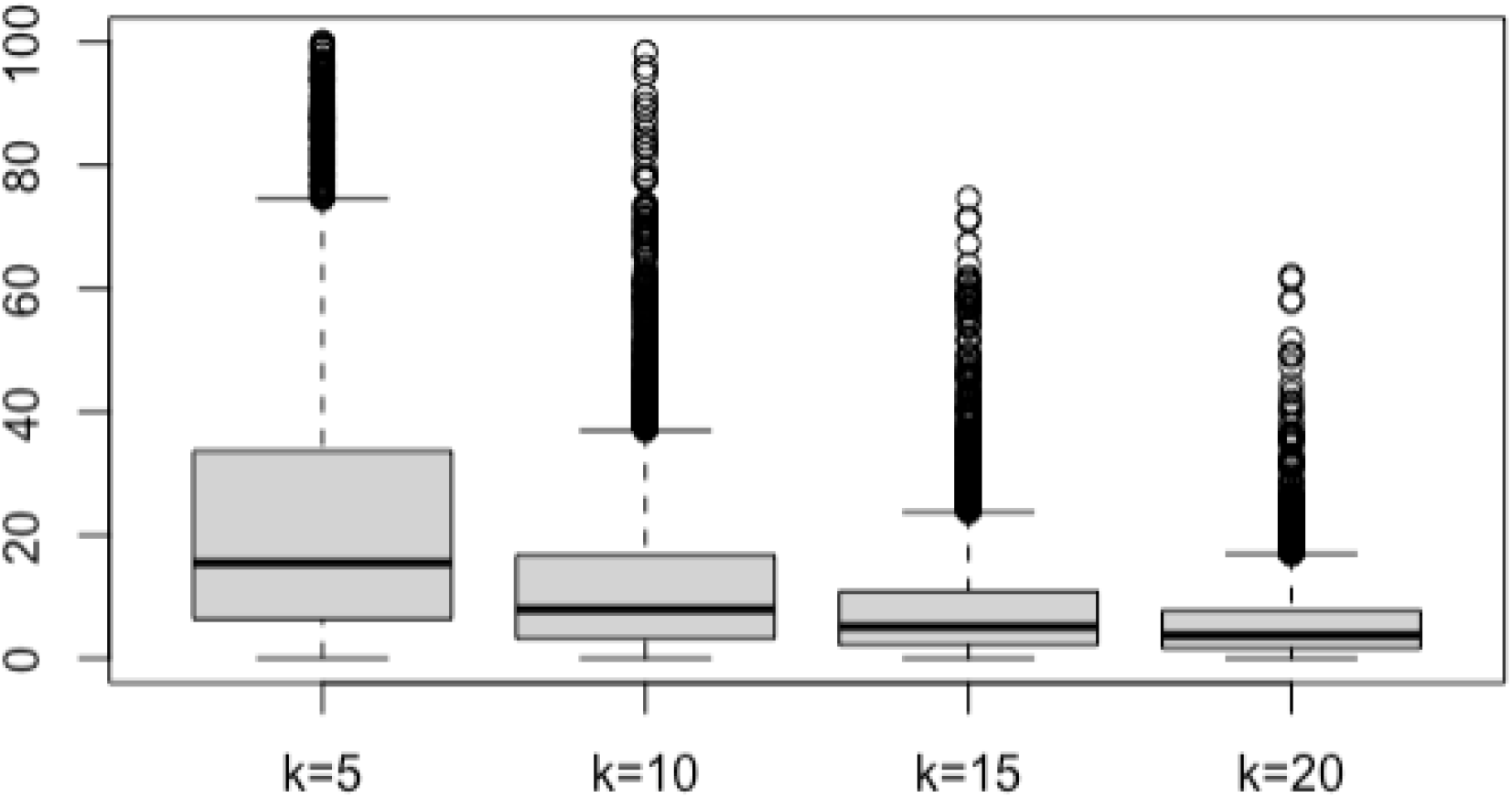
Boxplots of the centiMorgan lengths of spacings between within-chromosome repeats at 5, 10, 15 and 20 generation depths.

For comparison, note the overall mean length of a single ancestral segment at these depths is 16.6 cM, 9.09 cM, 6.25 cM and 4.76 cM. The ancestral genome corresponding in position to the non-ancestral intervening material may comprise several ancestral lineages from crossovers at a deeper generation than the original formation of the repeat (Figure 11). This is one reason why the consideration of the process of changes in ancestral lineage across a chromosome (Section 3.2) becomes complex.

### 4.3 Example of a chromosome ancestry

Figure 14 shows an example ancestry of a 1-Morgan chromosome. It is based on one of the 3000 chromosomes in the example simulation, but is perhaps atypical in that there are few events in the early generations, leading to just three simple segments of ancestry at generation 8. There is then a burst of activity, leading to 15 segments including 5 lineage repeats by generation 11. Thereafter repeats are lost, until there are 17 single segments at generation 14. Although generation 11 is extreme, and the reason for this choice of example, there is nothing outside the probability model in this example. Any exact series of events, viewed *a posteriori*, has low *a priori* probability.

**Figure 14:**
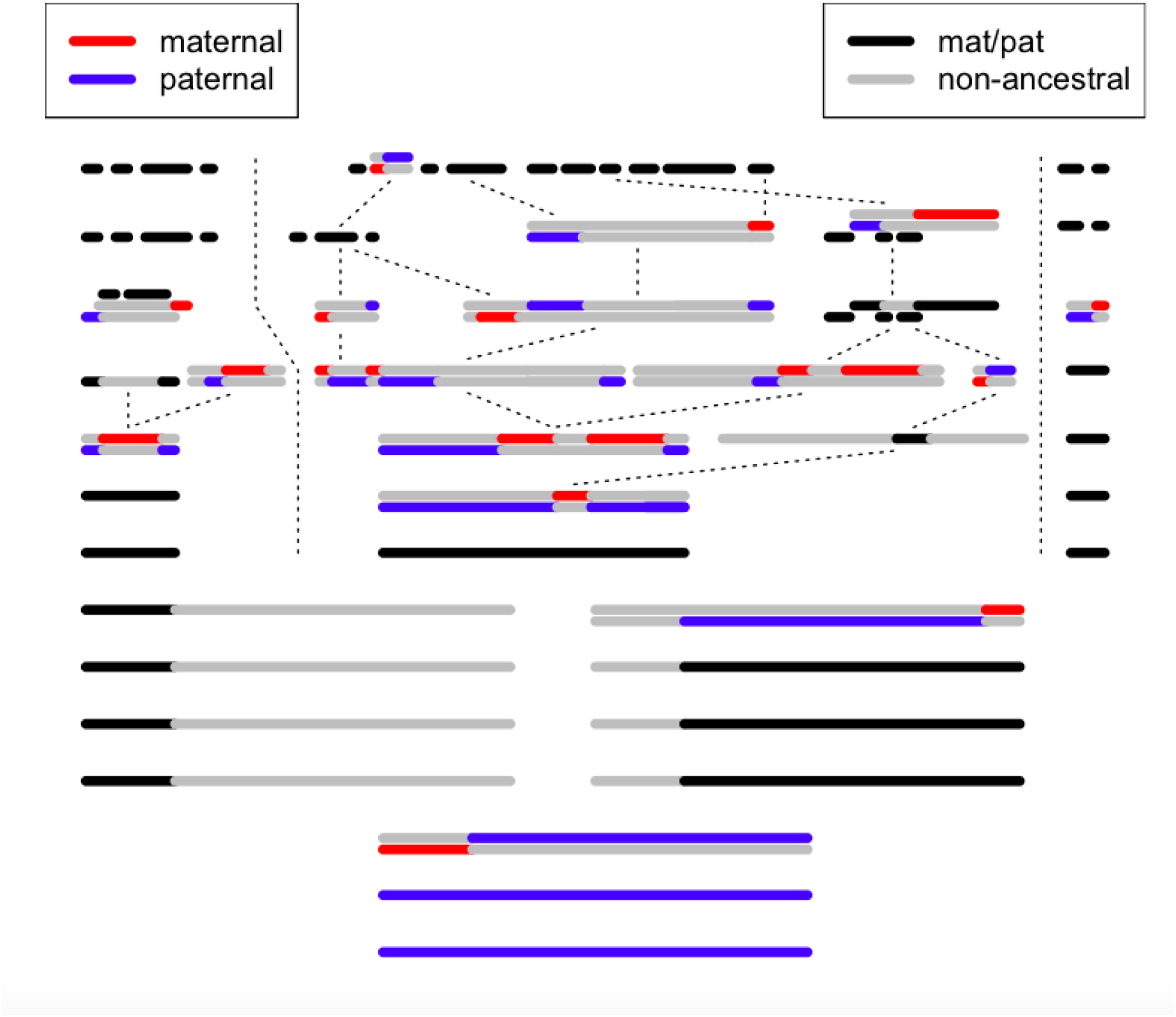
Example: the ancestry of a chromosome. See text for details.

### 4.4 The formation and loss of within-chromosome repeats

We now consider in more detail the formation and loss of within-chromosome repeats. The results are primarily empirical, using our sample of 3000 1-Morgan chromosomes, but Appendix B provides some rationale for the results. A repeat is formed when more than 1 crossover occurs in an ancestral segment (Figure 11). Ignoring multiple crossovers, which occur rarely after the first few generations, Equation (B.1) of Appendix B.1 shows that the expected number of repeats formed at generation *k* is (*k* + 1)/(*k* + 2)^2^ per chromosome. This result assumes than *k* is not small, and is poor for the initial generations of ancestry when many 1-Morgan chromosomes remain intact. Appendix B.2 discusses the formation of repeats in intact chromosomes, providing an adjustment for the first few generations. Considering separately the chromosomes expected to have remained intact from those that have undergone crossovers, and then combining the results, provides a better approximation. Figure 15 shows on the left the total number of repeats formed at each generation, together with this approximate expectation: the fit is good. On the right is shown the complete realized counts of repeats formed and surviving at each generation.

**Figure 15:**
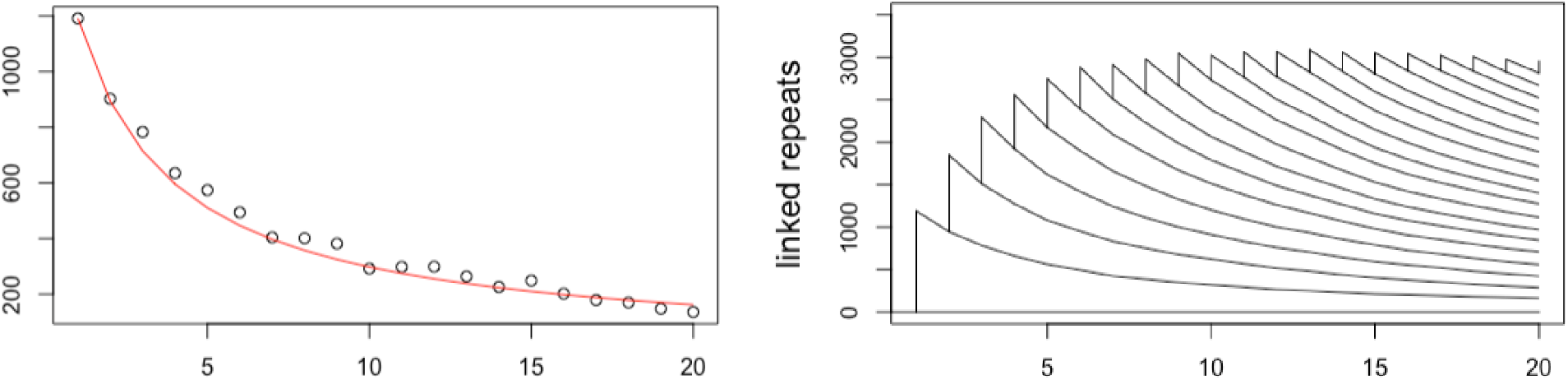
Formation and survival of repeats over 20 generations in 3000 1-Morgan chromosomes. Left: count of repeats formed each generation, with the theoretical approximation of the expectation in red. Right: realized totals of repeats formed and surviving.

Segments that form repeats undergo two (or more) crossovers, and hence are longer than average. Equation (B.2) gives the length of such an ancestral segment at generation *k* as (3*k* + 4)/((*k* + 1)(*k* + 2)), or approximately three times the average length. The formation of a single repeat by two crossovers divides the segment into two (repeated) ancestral segments and an intervening non-ancestral *spacing* segment (Figure 11). Conditional on this 2-crossover event, the positions are a uniform sample on the segment length, so each of the two ancestral segments and the intervening spacing have expected length *μ*_*k*_ = (3*k* + 4)/(3(*k* + 1)(*k* + 2)), or very similar to a random ancestral segment at this generation; note 1/(*k* + 2) *< μ*_*k*_ *<* 1/(*k* + 1). The exact probability distribution of the spacing length is unclear. Considering the 3000 realized chromosomes, and the repeats formed each generation, it appears to be approximately exponential. This may not be surprising, since, at the generation of formation of the repeat, this intervening length is simply an ancestral segment in the homologous chromosome of the parent (Figure 11).

Subsequent crossover events within the linked ancestral segments are not of great interest. Typically, they simply shorten the repeated segments. An exception would be if both ancestral segments were simultaneously split, with or without recombination in the non-ancestral segment. In this case both parental chromosomes would contain a repeat: however these are events of low probability. Of greater importance are recombination events within the spacing segment, since this is what causes the loss of the repeat (Figure 11). Note first that this is a completely independent process from crossovers in ancestral segments at this genomic location: that ancestry has diverged and will be in a different ancestral lineage. Nor does that intervening non-ancestral genome relate to any other chromosome under consideration. Second note that this is a recombination process (odd number of crossovers): an even number of crossovers in any meiosis in the non ancestral material will not affect the ancestry of the genome in any way. Appendix B.3 considered the loss of repeats. Equation B.3 approximates the 1-generation probability of survival of a random repeat as (1 + *μ*)/(1 + 2*μ*), where *μ* is the current mean of the spacing length distribution.

While the distribution of lengths of spacings in repeats is complex (Appendix B.3), some interesting findings are shown in the empirical set of 3000 chromosomes. Figure 16 shows mean lengths (in cM) of the spacing intervals of repeats. On the left are shown the means for repeats formed at generation *k*_1_ and lost at *k*_2_. Each line is for a given *k*_1_ *k*_1_ = 1, …, 19, and the x-axis gives *k*_2_, *k*_2_ = (*k*_1_ + 1), …, 20. It is not surprising that the means decline with *k*_2_. As shown in Section B.3 the distribution of spacing lengths is attenuated each generation through the loss of repeats with larger spacings. More surprising is the apparent independence of *k*_1_ in the mean spacing lengths for repeats lost at each given *k*_2_. Equation (B.4) provides some rationale for this result, but a larger simulation would be needed to study this more fully. On the right of Figure 16 is shown the overall mean spacing lengths for repeats formed at each generation (blue), together with the fitted curve from Equation (B.2) (black). Also shown in red is the mean for these same spacings for repeats surviving at generation 20. The theory provides a reasonable fit, and again, the mean for repeats surviving at generation 20 is largely independent of when they were formed, although appears to be slightly larger for the more recently formed repeats.

**Figure 16:**
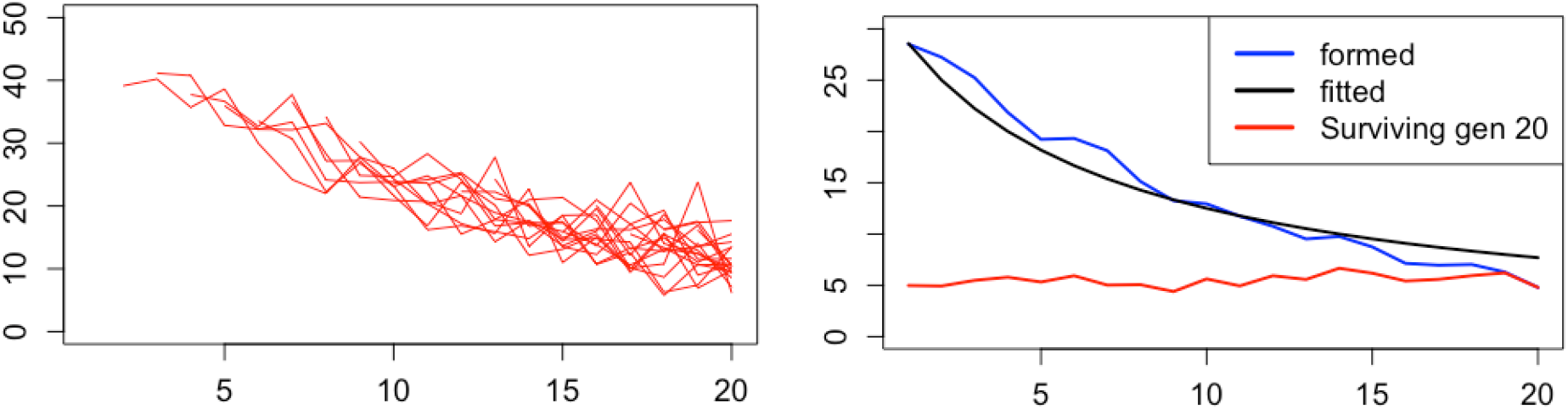
Mean lengths of spacings formed at each generation. Left: spacings lost at a given generation; Right: All spacings surviving to a given generation. See text for details.

The derivations in Appendix B assume not only that the distribution of ancestral segment lengths is exponential (which is broadly reasonable), but also that the spacing length distribution in surviving repeats is exponential, which is far from clear. However, Figure 17 shows that the approximation is close for the vast majority of spacings. Shown are qq-plots of spacing lengths (in cM) against an exponential distribution with the same mean. On the left is shown all spacings for the 2173 repeats formed in generations 1 to 4, and surviving at generation 5. (mean 18.81cM). The exponential distribution fits well except in the extreme 1% upper tail where the spacings are bounded by the chromosome lengths of 1 Morgan (100 cM): only 20 spacings are greater than 80 cM. On the right is shown the qq-plot of spacings for all 2816 repeats formed in generations 1 to 19, and surviving at generation 20 (mean 5.53cM). Again the exponential distribution fits very well except in the 2% upper tail, where spacings longer than expected under an exponential distribution survive. There are 60 spacings greater than 22cM, with 52.7 predicted, but there are 8 greater than 40cM with only 2 expected. Note that thee two sets of spacings represented in Figure 17 are the same as shown in the first (left) and last (right) boxplots of Figure 13.

**Figure 17:**
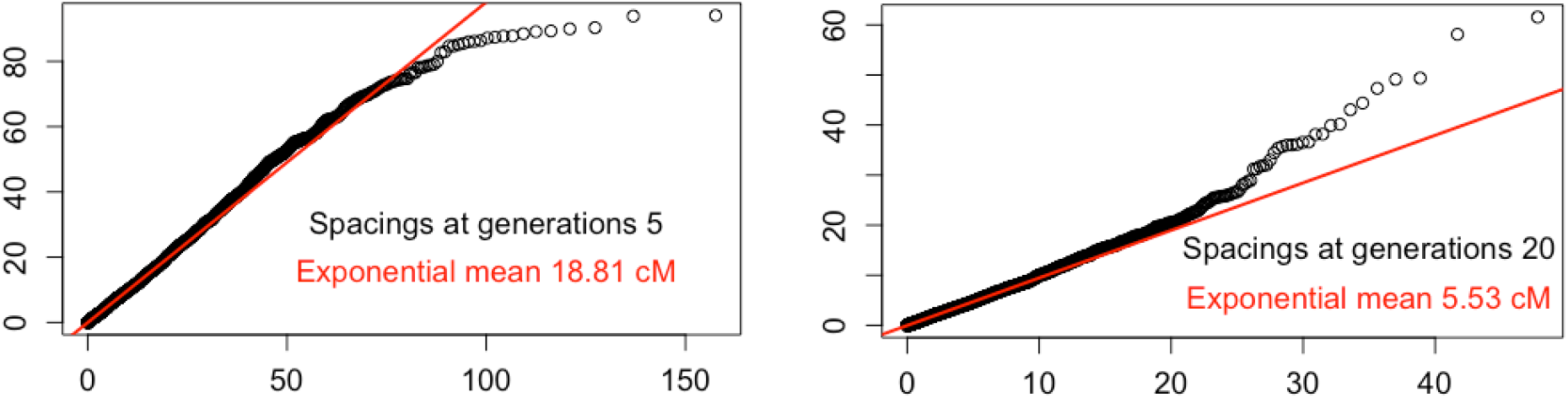
Lengths of spacings. QQ-plots of all spacings formed at generation 1 to (*k* − 1)) and surviving at generation *k*. Left *k* = 5. Right: *k* = 19.

## 5 Ancestors and genetic ancestors in a finite population

The above discussion has ignored coancestry, in effect assuming an infinite population. Once segments of ancestry diverge, they remain diverged, and each ancestral lineage references a different genome. In this section we provide a preliminary discussion of the effect of coancestry in a finite population on the counts of ancestors and genetic ancestors.

### 5.1 Haploid ancestry and descent in a finite population

To introduce the framework, we consider first a haploid genome. Suppose the effective population size is *M* and at time *k* = 0, 1, 2, 3, … back from the present we are considering the ancestry of *A*_*k*_ lineages. Conversely from a haploid genome *k* generations ago, we consider the number descendants at subsequent generations (*j* ≤ *k*). Each genome at *j* has a single parent at *j* + 1, and every genome *j* + 1 generations ago has an expected 1 offspring at generation *j*.

For this haploid case, unless *M* is very small, the descendant process is effectively a branching process with offspring count Poisson with mean 1: 𝒫*o*(1). The probability *q*_*t*_ of extinction within *t* generations is given by

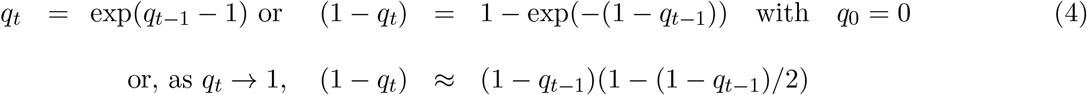

The probability of extinction increases from exp(−1) in one generation to over 0.91 over 20 generations. Conversely the expected number of descendant, conditional on survival, increases to 11.4 over 20 generations, and to 51.67 over 100 generations.

The ancestral process is more interesting; each haploid individual has a single parent, but a collection of haploids has a number of parents that depends on the coalescent process. Suppose that at *k* generations ago there are *A*_*k*_ ancestral lineages. Then at the preceding generation the probability an individual is not a parent of any of these lineages is 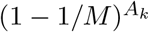, so the expected number of lineages at the preceding generation is

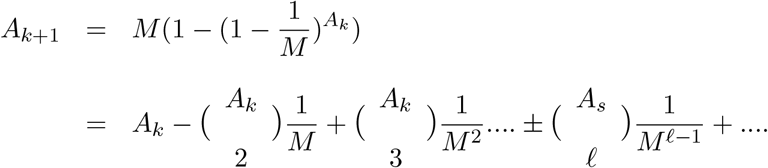

This formula shows as an inclusion-exclusion formula for the number of pairwise, three-way, …., *ℓ*-way coalescences. Figure 18 shows the number of ancestors of the current population over past generations: *A*_0_ = *M*, for varying *M* from 10^3^ to 10^7^. Proportionately the population size has little effect In every case the expected number of ancestral lineages at *k* = 100 generations ago is about 2% of the population size, corresponding to the expected approximately 50 descendants conditional on survival.

**Figure 18:**
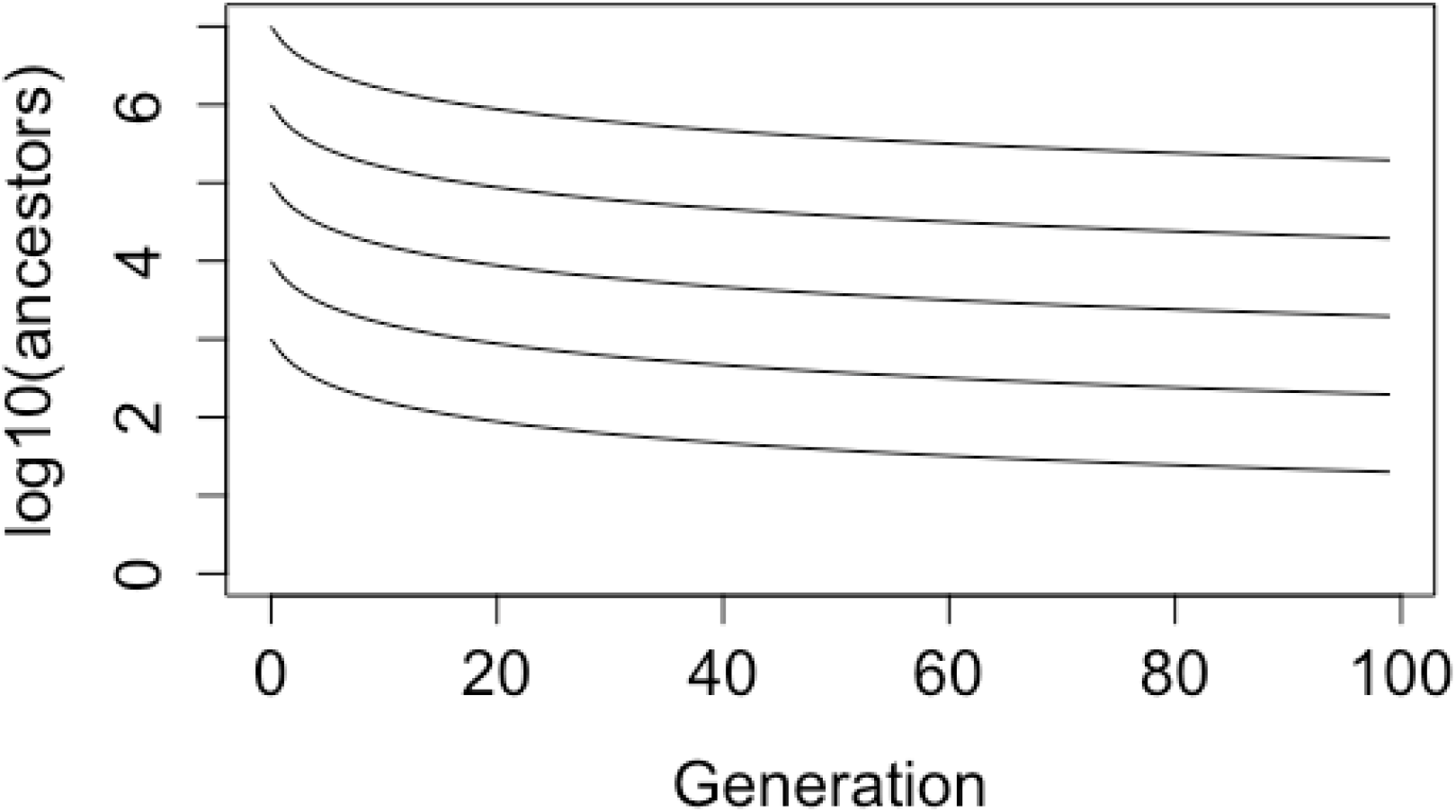
On a log scale, the number of ancestors of a haploid population, sizes *M* = 10^3^, 10^4^, …10^7^ at 1 to 100 generations ago.

### 5.2 Diploid descent and the ancestry

The picture for a diploid population is very different. First for an unbounded population, in which each diploid individual has a 𝒫*o*(2) number of offspring, the extinction probability equation (4) becomes

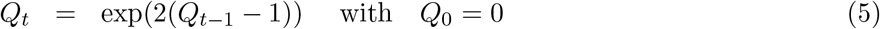

which increases rapidly from exp(−2) = 0.135 at *t* = 1 to the limiting value 0.203 by *t* = 8. Approximately 80% of diploid individuals have long-term descendants. In a finite random-mating diploid population of size 2*N* with *N* females and *N* males at each generation, every individual has two parents (one male, one female) and an expected 2 offspring. In a random-mating population, the probability of no descendants (diploid extinction) is little affected by population size, and equation (5) holds unless *N* is very small.

Note that the above haploid discussion (equation (4)) applies also to each single genome location in a diploid population. If each diploid has *X* ∼ 𝒫*o*(2) offspring, then the number of copies of a given parental DNA *W* is given by

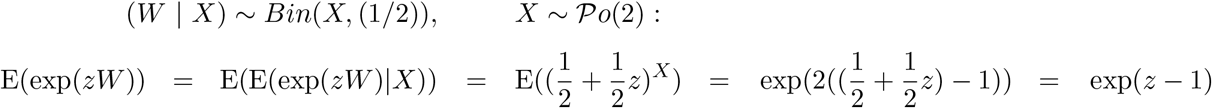

Or *W* ∼ 𝒫*o*(1). Thus the haploid model gives pointwise probabilities extinction of DNA in the diploid model and hence expected proportions of a genome extinct or surviving. It can also give probabilities for independently segregating and non-recombining units. For example, the probability that (in the absence of recombination) *C* chromosomes are all extinct over *t* generations is 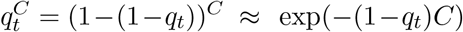. However, it does not give more detailed information on genome-wide ancestry.

Suppose a current individual has *A*_*k*_ ancestors, *k* generations ago (*A*_1_ = 2). Then the probability that a female [male] at generation (*k* + 1) is not a parent of any of these *A*_*k*_ individuals is 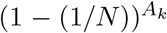 so that, combining males and females,

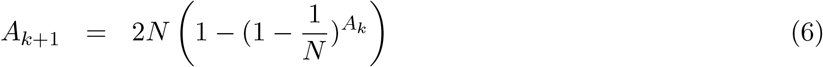

For *N* = 10^3^, 10^4^, 10^5^, 10^6^, Figure 19 shows, on a log10 scale, the number of ancestors of an individual both as a proportion of the population, and as a proportion of the nominal 2^*k*^ ancestors at generation *k*. Note that the limiting population proportion is not 1 (log_10_(1) = 0), but 0.797 (log_10_(0.797) = −0.0986), the limiting proportion of the population who have descendants.

**Figure 19:**
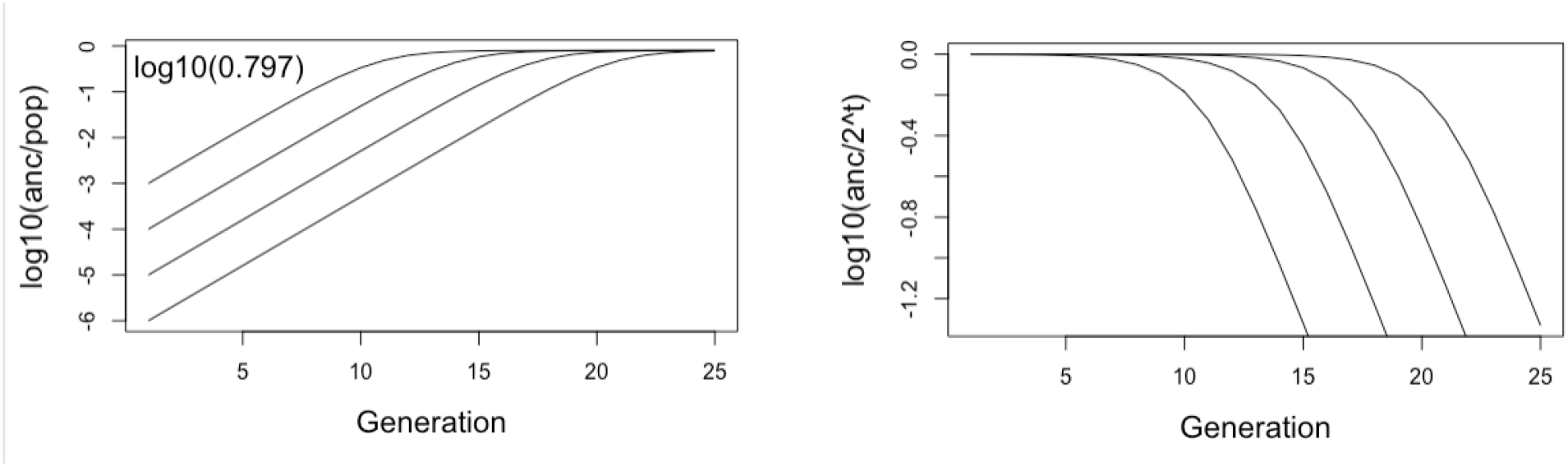
The number of ancestors of a diploid individual as a proportion of the population (left), and as a proportion of the potential number 2^*t*^ (right) both on a log_10_ scale.

The same formulae (6) can be applied to consider the ancestry of the entire population using *A*_0_ = 2*N*. The lines for proportion of the population than are ancestral to the current day are indistinguishable for the four population sizes 2*N* for *N* =!0^3^ to *N* ^7^, and within 8 generations the number of ancestors converges to the 79.7% of individuals who have long-term descendants, confirming the limiting extinction probability 0.203.

### 5.3 Genetic ancestry of a diploid in a finite population

We consider now the number of diploid genetic ancestors of a current haploid genome, for example the maternal genome of a current individual. We suppose that *k* generations ago there are *A*_*k*_ genetic ancestors, and consider the expected number at the preceding generation. Note that each genetic ancestor has either one or two genetic ancestor parents at the preceding generation. If one, then with equal chance it is either the male or female parent; if two, then both male and female parental genomes are represented at the preceding generation. There are three processes by which a genetic ancestor has both parental genomes represented, the first two being already considered in earlier Sections:

1. **Breaking of ancestral segments by crossovers:**. The total haploid ancestral genome length remains *L* Morgans at every generation, and there will be 𝒫*o*(*L*) crossovers in that ancestral genome. Each of the *A*_*s*_ ancestors has on average *L/A*_*s*_ length of genome, so the probability that genetic ancestor has only one parent that is a genetic ancestors is exp(−*L/A*_*s*_). As in the case of the infinite population, longer segments have higher probability of crossovers, so the fact that an ancestral segment is broken does not affect the probability of crossovers in the parents in the next earlier generation (Section 2.2).
2. **Recombination in spacing segments, breaking repeats:**. Both parental genomes are also represented if within chromosome repeats are broken by recombination. (Section 4.2 Figure 11). The within-chromosome formation (by more than one crossover in an ancestral segment) and subsequent loss is unaffected by the population size. Further, repeats form a decreasing proportion of the total ancestral segments, and are lost with decreasing probability (Appendiix B.3), so we will disregard repeat loss as a source of generating additional genetic ancestors. In effect, each cluster of tightly linked repeats is counted as a one ancestral segment.
3. **Independent merging of surviving ancestral lineages:**.If, for example, the mother’s, mother’s, mother’s mother is also the mother’s, father’s, father’s mother (the mother of the current individual is the child of half-first cousins) the lineages 000 and 110 both trace to the same diploid ancestor – a great-great-mother of the current focal individual (Figure 20). However, depending on what that individual segregates to his two offspring, it may not be the same (maternal/paternal) genome of that individual, and for more remote relationships surviving ancestral lineages referencing the same ancestor will likely be at genome locations on different chromosomes.

**Figure 20:**
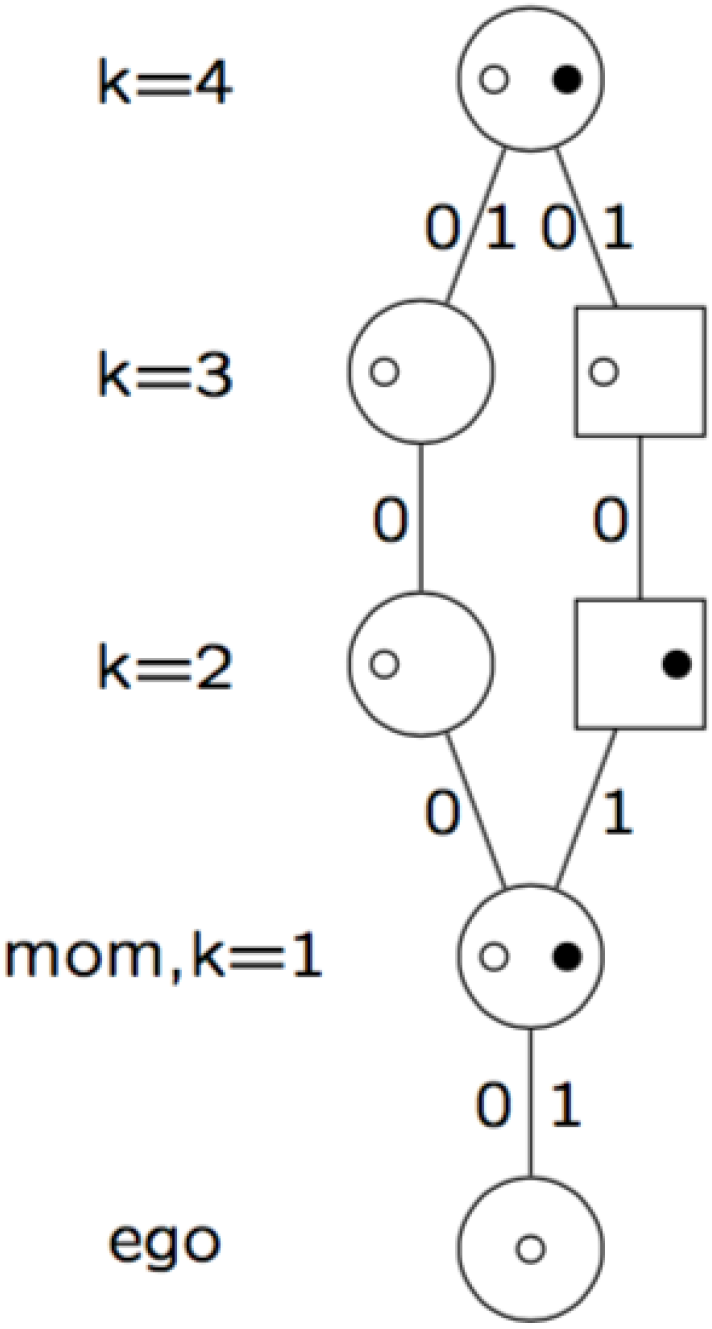
Coancestry over 4 generations.

In a finite random-mating population, ancestors may carry segments via different lineages. For the current discussion, we will assume these segments are not linked, and so independent in their parental origin. An ancestor at generation *k* carrying *S* segments via different ancestral paths will have only one genetic-ancestor parent with probability (1/2)^*S−*1^. We do not address possible matings of close relatives in this paper.

We now analyze in more detail the occurrence of multiple ancestral segments within an individual, due to matings of remote relatives. There are *C* +𝒫*o*(*kL*) segments at generation *k*. The segments are distributed over *C* chromosomes and most segments quickly disperse to a sparse subset of the total potential ancestral lineages (Figure 14). A few remain tightly linked (Figure 13) and a better approximation to the effective number of independently segregating ancestral segments is the expected number of genetic ancestors *G*_*k*_ in an unbounded population. From Section 2.3, *G*_*k*_ ∼ 𝒫*o*(*kL*) using the extinction formula of Donnelly (1983). If there are *A*_*k*_ diploid genetic ancestors, these *G*_*k*_ lineages are distributed among the *A*_*k*_ ancestors, each individual having at least 1. We assume the excess *D*_*k*_ = (*G*_*k*_ − *A*_*k*_) are distributed at random among the 2*A*_*s*_ ancestral genomes, so the number *S* of ancestral lineages carried by a genetic ancestor is approximately *S* ∼ 1 + 𝒫*o*(*D*_*k*_/2*A*_*k*_). (Note in an infinite population for large *k, G*_*k*_ ≈ *A*_*k*_, *D*_*k*_ ≈ 0, *S* ≈ 1.)

Combining the two independent processes of ancestral segment splitting, and merged lineages, the probability each of the *A*_*k*_ genetic ancestors has two genetic ancestor parents is

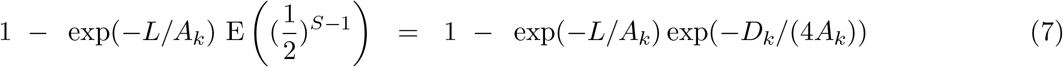

using the Poisson probability generating function to evaluate the expectation. The expected number of ancestors with both a male and female genetic parent is

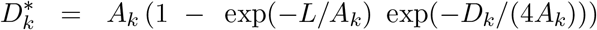

For the 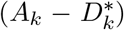 genetic ancestors who have a single genetic parent, there is a 50% chance it is the mother, so the number of (not necessarily distinct) mothers is 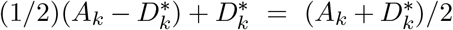, and likewise of fathers. Thus we may apply equation (6) to obtain

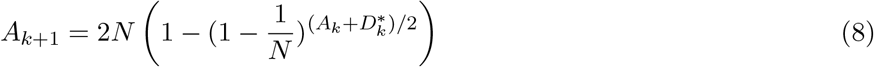

As a check, in an infinite population *D*_*k*_ = 0. Then when *k <* 5, *A*_*k*_ ≈ 2^*k*^ *< L*, and 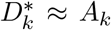.

All ancestors have two genetic ancestor parents. (Section 2.3). When *k* is large, *A*_*k*_ ≈ *kL* and 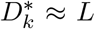, if *L* is not small. When the number of segments *kL* is several times the number of ancestors having any current descendants (either because *k* is large, or the population size small) *D*_*k*_ ≈ *G*_*k*_ ≈ *kL*. and again 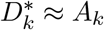 – each genetic ancestor carries several ancestral segments, so that both parents are very likely to be genetic ancestors. Note also that when 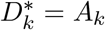, equation (8) reduces to equation (6). We note that we are considering here relatively recent history, say *k* ≤ 100, and not evolutionary time scales. Clearly the genome cannot be infinitely fragmented fragmented, and, in a finite population, ultimately identity by descent will occur for all individuals at every point in the genome.

Figure 21 shows the result of applying equation (8) for a range of orders of magnitude of population size, from 200 to 2 million, each generation consisting of an equal number *N* of males and females and also comparing with the infinite population result 2^*k−*1^(1 − exp(−(*k* − 1)*L/*2^*k−*1^))). (The one-generation offset occurs because we consider the diploid ancestors of a current haploid genome, or of a generation-1 diploid mother; see Donnelly (1983).) We see that at generation *k* ≤ 100 the line for a population of size 2 million is indistinguishable from the infinite-population case, and that for 2*N* = 200, 000 there is scarcely any effect. For a population of 20,000 there is some effect, with the rate of increase of ancestors per generation being 20 rather than 30, by *k* = 100. Only for a small population of 2*N* = 2000, is there a major effect within 100 generations, with a rate of increase about 4 per generation and total number of genetic ancestors just over 1000 at *k* = 100, which is about 62% of the total number of the 1600 diploid ancestors. In the very smallest populations 2*N* = 200, the number of genetic ancestors at 159 appears to have reached the limiting bound of the total number of ancestors, although the adequacy of the model and approximations is doubtful for such a small population. The level of pointwise inbreeding at 100 generations is 0.386 in a random mating population of 100 males and 100 females (Crow and Kimura (1970); P. 103 Eq. 3.11.6). The numbers produced by equation (8) should be taken as an upper bound on the expected number of genetic ancestors..

**Figure 21:**
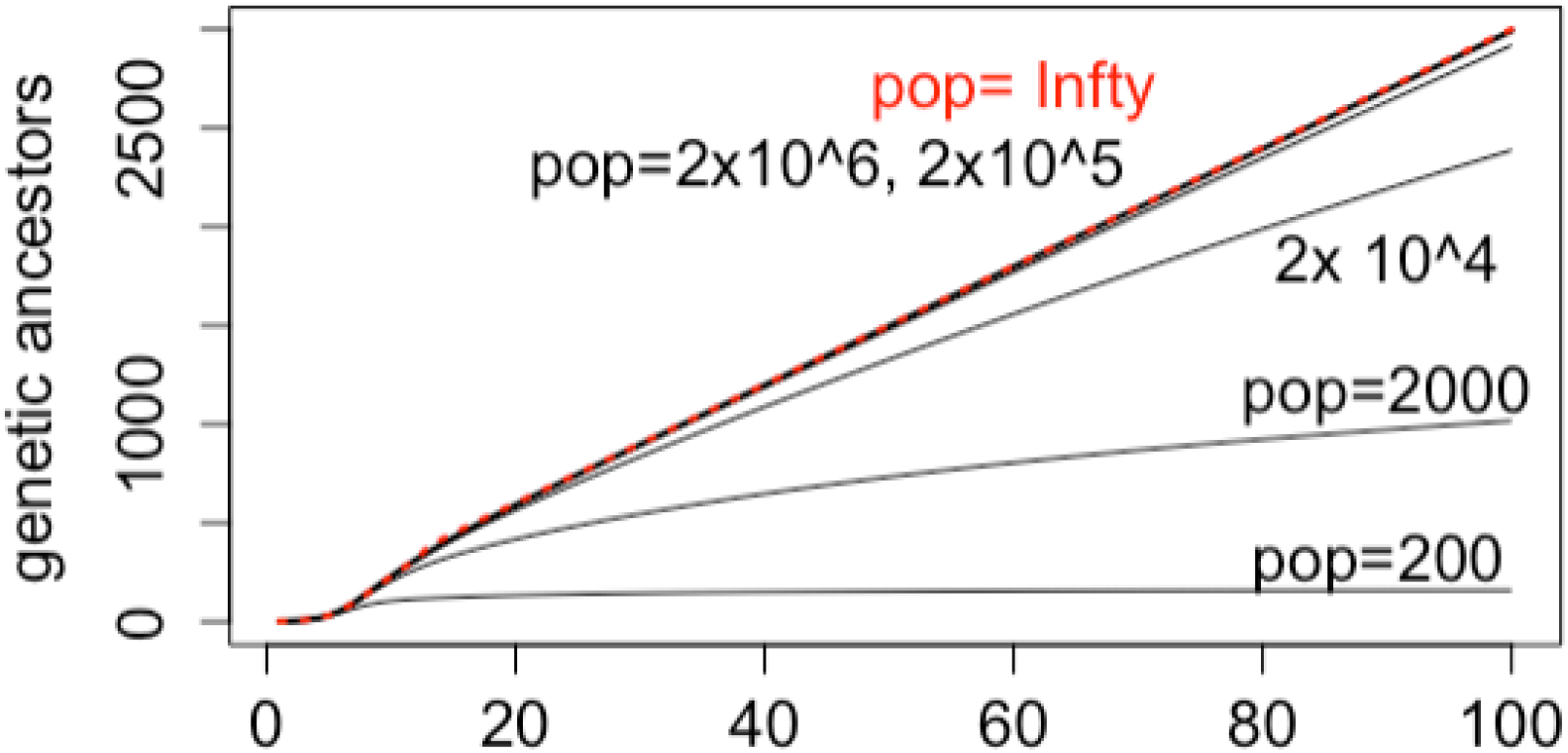
The number of genetic ancestors of a diploid individual over 100 generations, for population sizes 2 × 10^*m*^, for m=2,3,4,5,6, and in an unbounded population.

### 5.2 The effect of migration

Populations are not random mating, being subject to many forms of substructure. Here we consider a simple geographic stepping-stone model as a first view of the effect of population structure. We consider 961 communities each of size 1000 males and 1000 females, on a 31 × 31 toroidal grid, for a total population close to the 2 × 10^6^ considered above (Figure 22). At each generation, migration occurs at equal rates *m/*4 to adjacent communities in each of the four directions.

**Figure 22:**
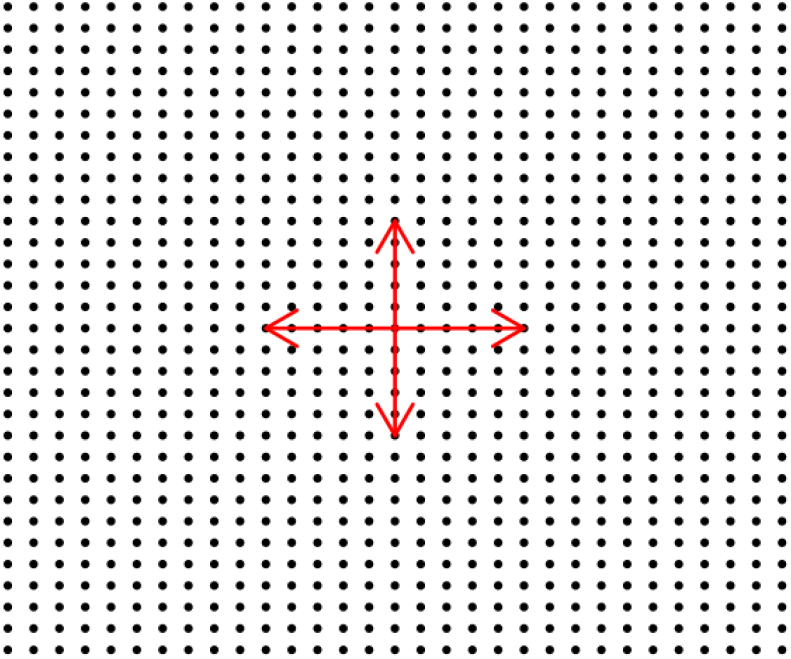
Toroidal 19 by 19 grid of 961 communities each of 2000 individuals

We apply the same formulae as above for the next-generation ancestors in each community, and then out migrate to the 4 neighbors at the specified rate. When the expected number of ancestors in a grid point is *x* ≤ 1, the number of parent ancestors in the same grid point is 2*x* and migration does not occur.

Figure 23 shows results for the number of diploid ancestors of a single individual. On the left, on a log_10_ scale are shown in red the total expected numbers under a high migration rate (total migration *m* = 4 × 0.025 = 10%: 0.025 in each of 4 directions for each individual at each generation) and in blue under a low migration rate (*m* = 4 × 0.0025 = 1%). The curve for random mating populations size 2 × 10^*n*^ for *n* = 3, 4, 5, 6 (see also Figure 19) are shown for comparison. For the higher migration rate (red) the total number of ancestors rises quickly, surpassing the number for a random mating population of size 20,000 and size 200,000 at about 15 and 25 generations respectively. For the low migration rate the number of ancestors increases more slowly, but still approaches the total number of ancestors in a population of almost 2 million (i.e. 79.7% of that total population) by 100 generations.

**Figure 23:**
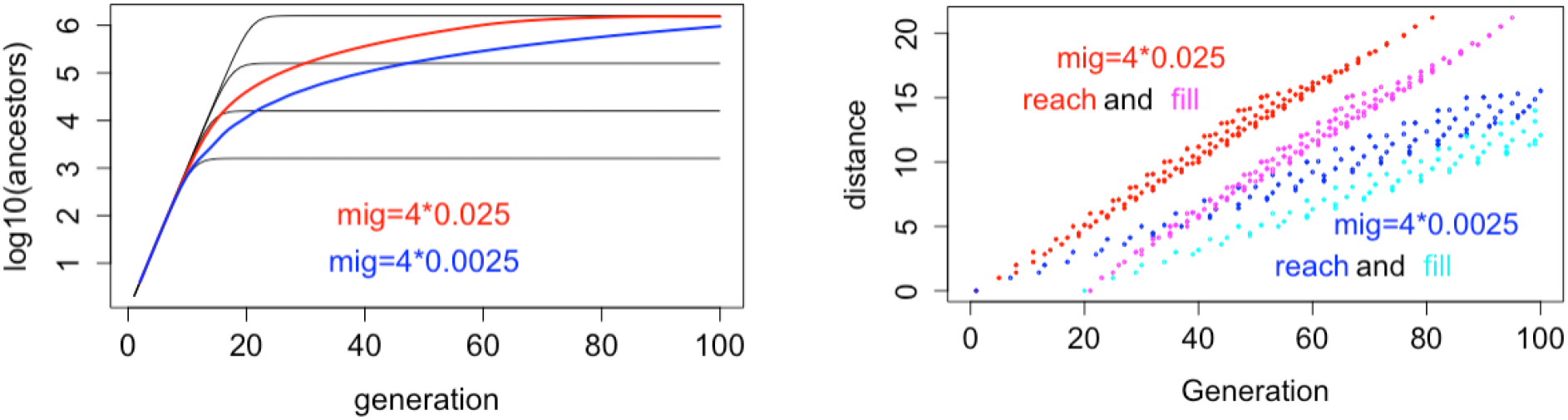
Left: The number of ancestors of a diploid individual over 100 generations under migration. Right: the distance of communities expected to be reached (red and blue) and filled (magenta and cyan) by a given generation.

The right-hand figure of Figure 23 explains these results. As soon as an ancestor reaches a community of 1000 males and 1000 females, the number of ancestors there increases very rapidly to the almost 80% of individuals having long-term descendants (Figure 19). The right figure of Figure 23 shows the generation that the expected number of ancestors in a community at a given distance exceeds 1, and the generation at which it exceeds 79.5% of that local population, where distance is the Euclidean distance of each community grid-point from the community of the focal individual. With a total 10% migration rate, the furthest communities (distance 22) are reached by 75 generations, and reach their limiting ancestral count within 15 generations later. With the smaller 1% migration rate, once communities are reached, there is a similar rapid increase to the limiting number of ancestors. However, only populations closer than distance ∼ 16 are reached in 100 generations, and only populations at distance less than ∼ 13 from the original individual have been reached long enough for the number of ancestors to have reached the local limit.

Computations for the expected counts of genetic ancestors under migration are more complex. Although a version of equation (7) may be implemented iteratively over generations to estimate the expected numbers of genetic ancestors in different communities, the accumulating geographic inhomogeneity of amounts of genome per ancestor *L/A*_*k*_ and of the distribution of excess segments of ancestry *D*_*k*_ among individuals in each community renders this at best a crude approximation. Further the small integer counts of migrating ancestors are not well represented by their expectations. The results shown in Figure 24 should be considered upper bounds. The left figure shows that with the higher migration rate (total 10%) the number of genetic ancestors crosses the line for a single population of 20,000 at about 25 generations, and thereafter follows only somewhat below the large-population value. With the lower (total 1%) migration rate, the count passes the count for a single population of 20,000 at about 60 generations. Note that a total of 20,000 individuals corresponds to a Euclidean distance about 3 around the focal point. The right figure of Figure 24 shows that greater than 1 expected genetic ancestor reaches communities of distance 6 within 100 generations for total migration rate 10%, while for migration rate 1% only communities within distance 4 are reached. These correspond to 37 and 15 communities or total population sizes of 74,000 and 30,000 respectively. Of course, unlike for total ancestors, the genetic ancestors do not ”fill” a community, but remain a small fraction of the total individuals. Also note the apparent irregularity of the points in Figure 24(right) is a result of the Euclidean distance metric on this small number of discrete communities on an integer grid.

**Figure 24:**
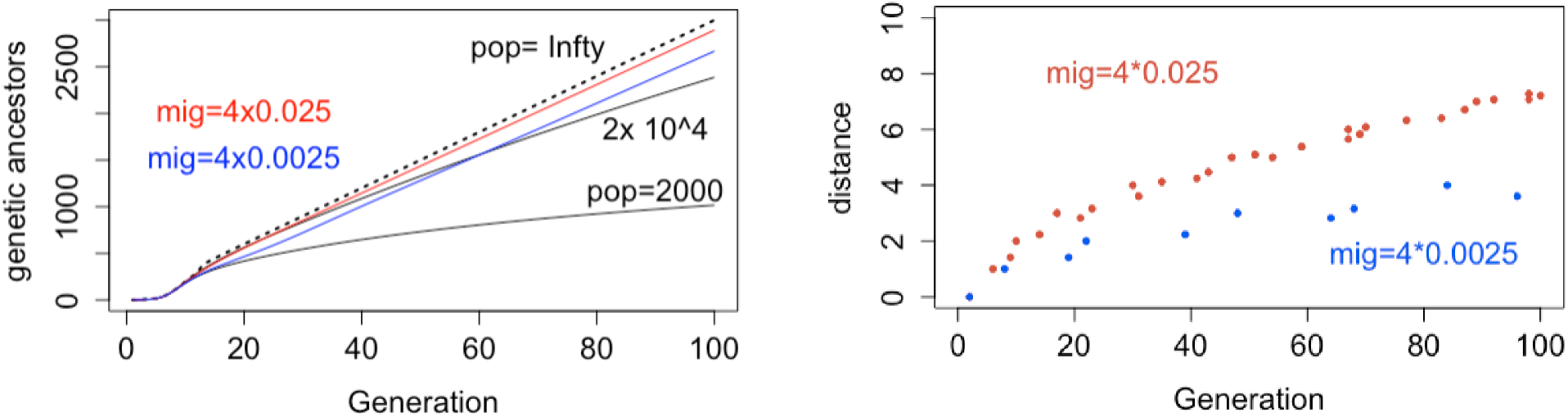
Left: The expected number of genetic ancestors of a diploid individual over 100 generations under migration. Right: The distance of communities reached by a given generation.

A simpler and more accurate computation is of the expected proportion of an individuals’s DNA that descends from ancestors within a given distance from the focal individual. This expectation over the genome is obtained from probabilities at a single genetic locus, with DNA at each locus in the genome deriving from exactly one parent at each generation (Section 5.1). The results are shown in Figure 25 for generations 20, 60 and 100 for the higher 10% migration rate, and at 100 generations for the 1% migration rate. For the 10% migration rate, at 20 generations almost 25% of the ancestral DNA remains in the focal community, while almost all is within distance 3; by 60 and 100 generations only a small percentage remains in the community of origin, but a very high percentage is within distance 6 or 7. For the 1% total migration rate, dispersion of ancestral DNA at 100 generations is less than for the 10% migration rate at 20 generations. Over 42% remains in the community of origin, while barely any reaches even a distance of 3. (Again the Euclidean distance metric accounts for the uneven discrete set of distances.)

**Figure 25:**
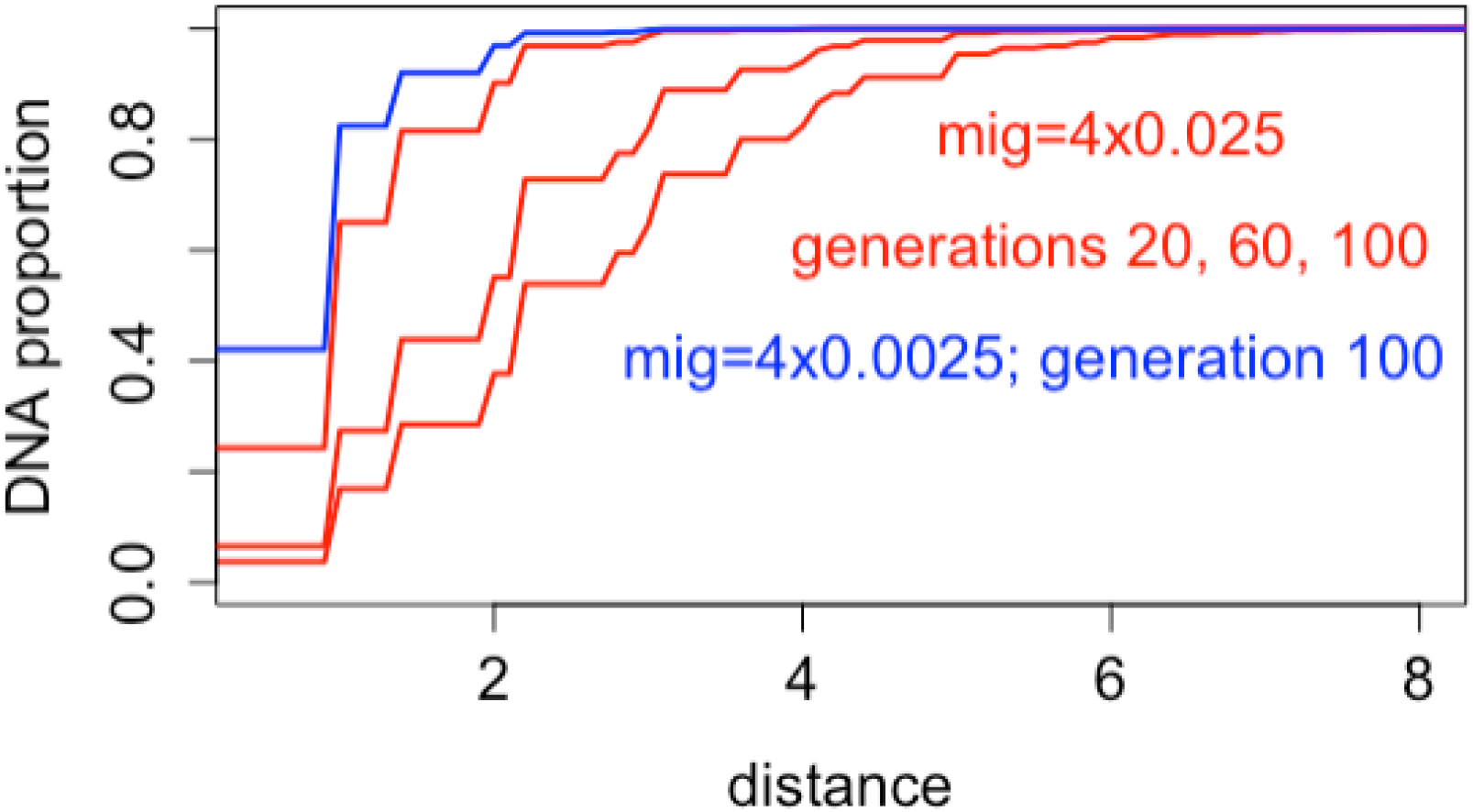
The expected proportion of the genetic ancestry of a diploid individual that is within a given distance of the origin at up to 100 generations. See text for details.

## 6 Discussion

This paper has explored the patterns of ancestry of a current genome, focusing on the numbers of genetic ancestors at past generations, the clustering of represented ancestral lineages in the totality of potential lineages, and the patterns of surviving ancestral genome segments. Much of this work originates with Donnelly (1983), who provides not only key results but much insight and discussion. However, in the case of ancestry, that paper focuses on survival of genome of each single lineage. In contrast, here we have considered the diversity of lineages represented in a current genome.

The exponential distribution is of course key to these analyses, as the genetic distance to the next crossover event under the Haldane (1919) model of no genetic interference. However, shape of the exponential distribution also underlies some of the more surprising results. It gives rise to numerous very small values, and a few very large ones, providing non-negligible probabilities of multiple crossovers within a single ancestral segment even at distant generations into the past. It also appears unexpectedly, as a remarkably accurate approximation to the distribution to many of the statistics considered. Especially in Section 4.4, there seems to be no good reason for the distribution of the genetic lengths of accumulated surviving spacings between repeated ancestral lineages to be exponential. Yet the qq-plots based on almost 3000 values show that the distribution diverges from exponential only in the extreme tail.

The long-term process of formation and loss of these repeat ancestral lineages merits further study. Over the time period we consider, formation of repeats is closely balanced by loss, and the number surviving remains remarkably constant. As time *k* into the past progresses, ancestral segments become smaller at rate *k*^−1^. However, the number of segments increases linearly in *k*, so the expected number of newly formed repeats also decreases only as *k*^−1^. Moreover, surviving repeats are more tightly linked, and the loss of repeats also decreases. The apparent distribution of spacings in repeats suggests this process also follows a *k*^−1^ rate. Ultimately, in reality this process must break down, since the genome is not infinitely divisible. However, the extent to which genomes carry such repeats from their distant ancestors remains a topic to be explored.

Our basic study was of ancestry in an unbounded population, but in reality populations are finite. Since genetic ancestors are few, in a large random-mating population of order size 10^5^, the constraint of population size has little impact on the number of genetic ancestors over 100 generations. The study of a large structured finite population with stepping-stone migration shows that while our ancestors are many and everywhere, our genetic ancestors are few and local. This is not unexpected: by the time (tracing backwards) an ancestor reaches a more remote population, the probability they are a genetic ancestor is minute. However, the possibility of long-range migration, even at low probability, could overcome this. Back five generations, all our ancestors are typically genetic ancestors: a distant migrant would establish a new local area of genetic ancestry. This would be an interesting process to explore, as would other classical migration models, such as an Island Model (Crow and Kimura (1970); Section 9.2).

Our study of finite populations has ignored inbreeding. That is we allow for ancestral segments to arrive in the same ancestral genome via different lineages, but not to coalesce at the same point in the genome. Clearly, even over only 100 generations, this is an approximation in a small population, Even in a large population, the number of genetic ancestors cannot increase linearly indefinitely. Our ancestors are approximately 80% of the total population, and even in small populations the event of having long-term descendants is determined within 5 or 6 generations, a period over which almost all ancestors are genetic ancestors. In small populations (for example, size 200), it appears that the number of genetic ancestors approaches the number of ancestors. Is this ultimately true in larger populations? Do we have ancestors who are not genetic ancestors? Clearly there is non-zero probability of this event, but the probability is of interest. In a large but finite population, the reality that the genome is not infinitely divisible may ultimately bound the probability. In a small population, the probability may be very small.

By making the approximation that the ancestral segments do not coalesce, even when they arrive in the same ancestral genome via different lineages, we essentially do not allow for the mating of close relatives. In the study of pedigreed populations of severely endangered species this is clearly inadequate. Pointwise inbreeding levels may be high (Thompson, 2023), and the number of ancestors quite limited. A study of the process of ancestral genome segments under mating of relatives, or within a defined pedigree is an area beyond the scope of this paper.

The paper arose from an interest in exploring patterns of ancestry in an individual recently descended from a large population. It makes no great claims of impact, either in analysis of recent human evolution, or in conservation genetics. However the results seem to be of some interest, and give rise to many further questions to be explored.

## Acknowledgment

I am grateful to Dr. Joe Felsenstein and Dr. Mary Kuhner for many helpful discussions and comments.

## Statements and declarations

No funding was received to assist with the preparation of this manuscript. The author has no relevant financial or non-financial interests to disclose.

## Appendices

### A Transition process of ancestral lineages across a chromosome

#### A.1 A small example: The case *k* = 3

**Figure A1:**
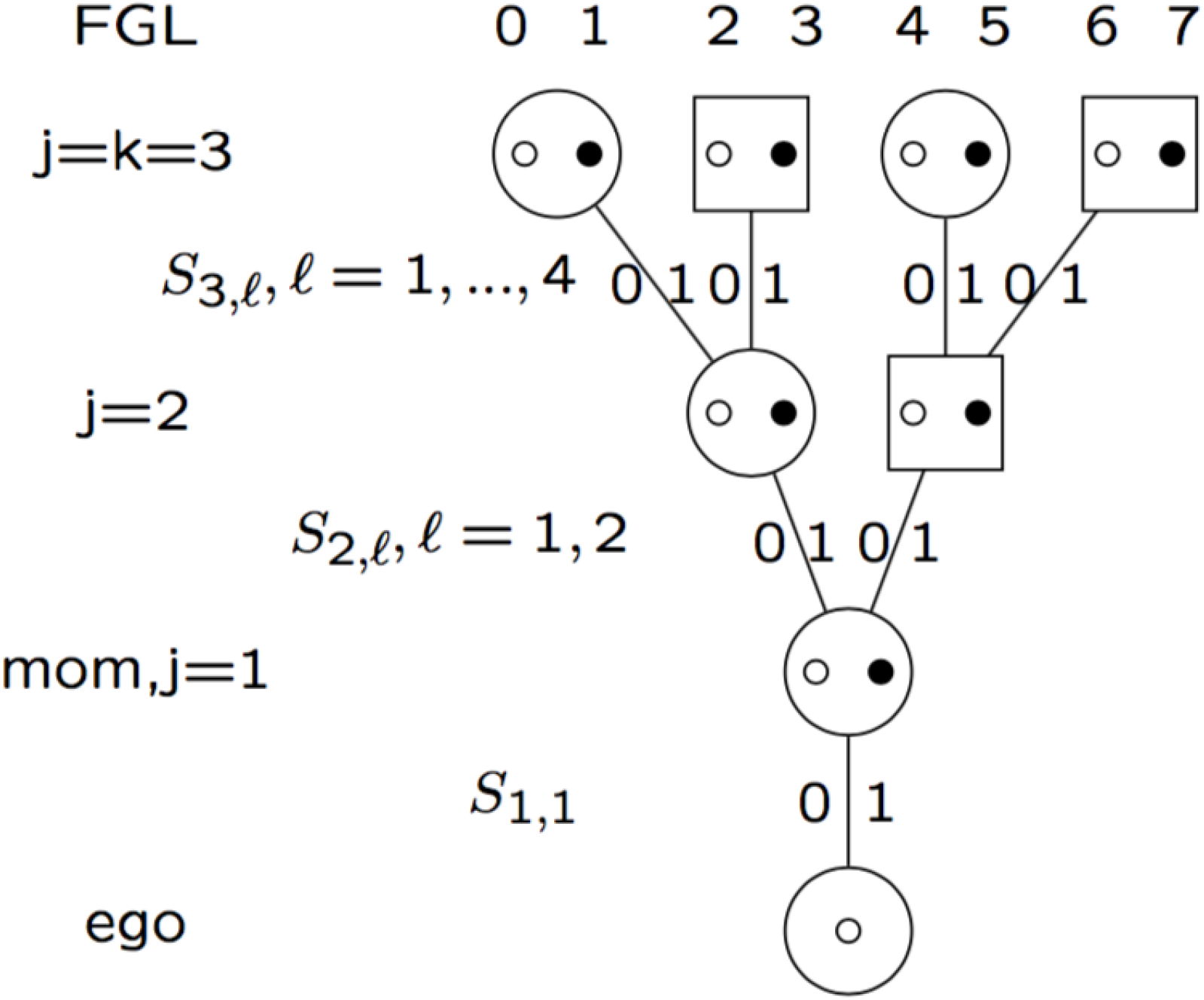
The meiosis labeling and structure for the case *k* = 3

To clarify the above we work in detail through the case *k* = 3: There are a total of 7 meioses, and 8 FGL is shown in Figure A.1. Each FGL is determined by the value of *k* meioses, one at each level. Thus, for example when *k* = 3, the maternal lineage *S*_1,1_ = *S*_2,1_ = *S*_3,1_ = 0 results in FGL 0, while *S*_1,1_ = *S*_2,1_ = 0, *S*_3,1_ = 1 results in FGL 1. The fully paternal lineage *S*_1,1_ = *S*_2,2_ = *S*_3,4_ = 1 results in FGL 7. There are 128 meiosis patterns, 8 FGL, each corresponding to 16 = 2^4^ meiosis patterns.

Suppose the lineage at the start of the chromosome has FGL 0: *S*_1,1_ = *S*_2,1_ = *S*_3,1_ = 0, with the remaining four meioses each being 0 or 1 independently with probability 1/2. Then there is probability 1/3 that the first change of lineage will be to FGL set. {1}, {2, 3} and {5, 6, 7, 8}, with equal probability to each member of the subset and hence probabilities (1/3, 1/6, 1/6, 1/12, 1/12, 1/12, 1/12) for FGL 1,2,3,4,5,6,7. Results for other initial FGL are given by symmetry resulting in the following one-step transition matrix among FGL 0 to 7:

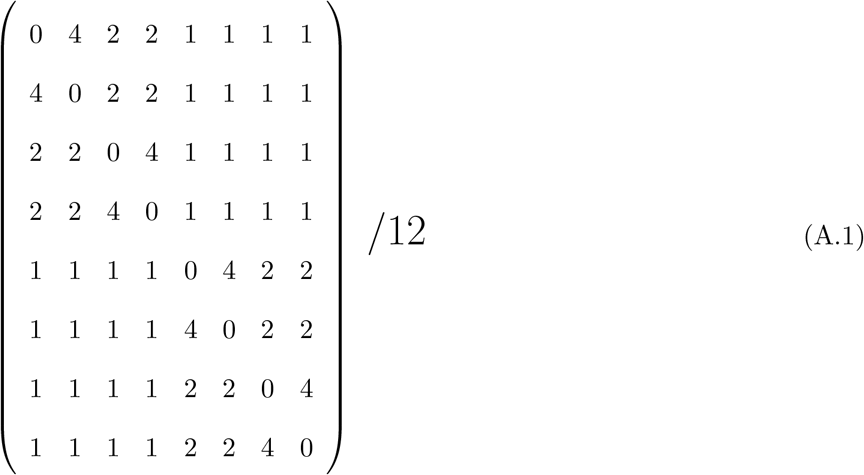

#### A.2 A key result

##### Lemma

**Proof 1:** Since the meioses switch independently, only the (*k* + 1) meioses are relevant, and at every change the index meiosis is the one to switch with probability 1/(k+1). In order for the index meiosis to be switched when the first of the other *k* meioses switches it must switch

(first, but not also second) or (three times but not four) or …..((2*ℓ* + 1) times but not also (2*ℓ* times) ….

with probability

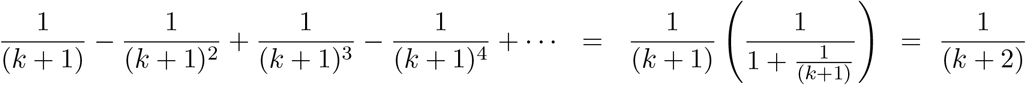

**Proof 2:** The distance to the switch of the first of the set of *k* meioses is the minimum of *k* independent standard (mean 1) exponential random variables, with is an exponential mean 1*/k* with probability density function *k* exp(−*kt*) on 0 *< t <* ∞. The probability that the index meiosis is in changed state at distance *t*, is the recombination probability (1 − exp(−2*t*))/2 of equation (1) above. So the probability the index meiosis is in changed state is

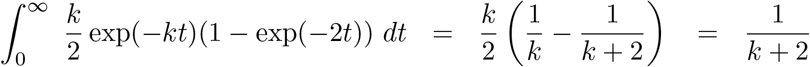

This result provides the probability 1/(*k* + 2) that any meiosis that was in the immediately precediing lineage, but not in the current lineage of *k* meioses is in changed state at the point on the chromosome at which the lineage changes again.

#### A.3 Two-step-transitions: the case *k* = 3

We now use the lemma of Appendix A.2 to obtain the two-step transition probabilities for the case *k* = 3. Suppose the current FGL step is 0 (*S*_1,1_ = *S*_2,1_ = *S*_3,1_ = 0), and consider the preceding state being each of *X* = 1, 2, 3, 4, 5, 6, 7 in turn, with no additional information. Note that all other two-step transitions will be given by symmetry.

(a) If *X* = 1, then *S*_3,1_ switched from 1 to 0 (Figure A.1), and there is no information on the other 4 meioses, and the next-step transition probabilities are as for the 1-step transitions above.

(b) If *X* = 2, then *S*_2,2_ switched from 1 to 0 (Figure A.1). The next switch from FGL=0 will be to FGL=1 with probability 1/3 (*S*_3,1_ = 1), and to FGL set {4, 5, 6, 7} with total probability 1/3 (*S*_1,1_ = 1), with equal probability to each, since there remains no information about *S*_3,3_ and *S*_3,4_. If the switch is back to FGL set {2, 3} (*S*_2,2_ = 1; probability 1/3), then the resulting FGL will be 3 if *S*_3,2_ is in changed state (probability 1/(3+2) = 1/5) and will be FGL 2 otherwise (probability 4/5).

The case *X* = 3 is given by symmetry.

(c) If *X* = 4 (with cases *X* = 5, 6, 7 given by symmetry), then *S*_1,1_ switched from 1 to 0 to arrive at FGL=0. In this case there is no information about *S*_2,2_ and *S*_3,2_, so the next state change is to FGL=1 with probability 1/3. and to FGL=2 and FGL=3 each with the one-step probability 1/6. The remaining probability 1/3, from reversing the switch of *S*_1,1_, is divided between states {4, 5, 6, 7} as follows. If *S*_2,2_ is in changed state (probability 1/5) the move will be to FGL=6 or FGL=7, split equally between them as there is no information about *S*_3,4_. If *S*_2,2_ is unchanged (probability 4/5), then if *S*_3.3_ is unchanged (probability 4/5) the state returns to FGL=4, while if *S*_3,3_ is changed (probability 1/5) the resulting state is FGL=5. Note all the above results are dependent on the basic property that meioses switch independently and at equal rates.

The transition probabilities above may be summarized in the following matrix, where *X*_−1_, *X*_0_, and *X*_+1_ indicate a previous, current, and next lineage across the chromosome:

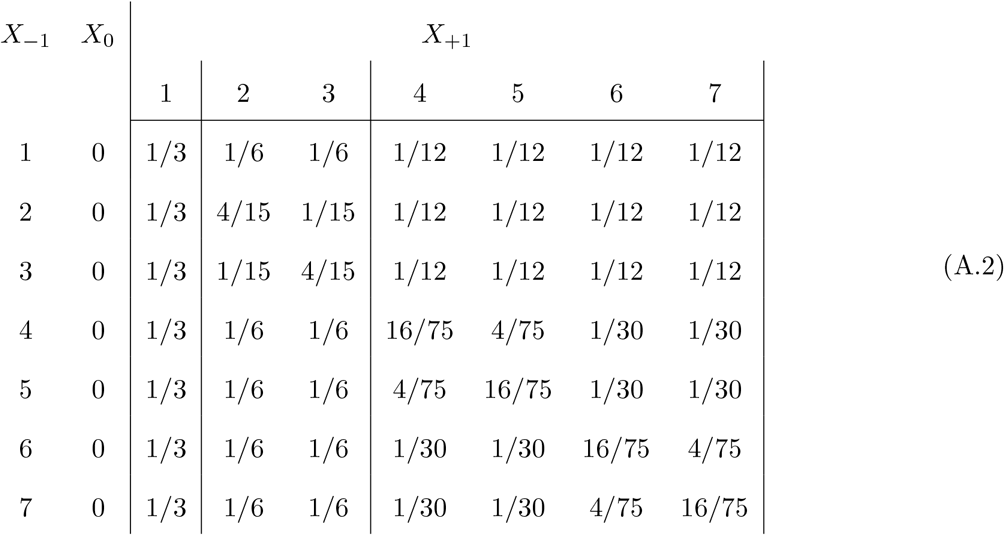

All other elements of the 2-step transition matrix are given by symmetry; that is by the exchangeability of pairs of FGL that are connected at the same depth-level *j, j* = 1, 2, 3.

### B Details of within-chromosome repeat computations

#### B.1 Formation and lengths of within-chromosome repeats

We now consider more mathematically the formation (See Figure 11) and retention of within-chromosome *repeats* of ancestral lineages, separated by non-ancestral *spacing* segments. The analysis here involves a number of approximations, but provides a framework for understanding the process and results. Specifically we ignore the possibility of more than two crossovers in one ancestral segment in one meiosis, which simultaneously creates more than one repeat: one repeat in each of the two parental chromosomes if there are 3 crossovers, and more than one repeat in either parental chromosome if there are more than 3 crossovers. Even for small generation depths *k* these are relatively rare events.

We also ignore dependence over successive generations. A segment than has been broken by two crossovers has longer than average length, but the fact that it now consists of two ancestral segments with intervening non-ancestral genome, mean that these segments no longer have the same overall marginal length distribution at the next generation. This will affect probabilities of their subsequent ancestry. The overall effect of ignoring this dependence appears to be slight. We will also ignore the truncation of segments by chromosome breaks: this has little effect for larger *k*, but does permit the existence of exceptionally long ancestral segments which are more likely to contain multiple crossover points and hence repeats. Since, for larger *k*, we are considering events on much less than a 1-Morgan scale, our integrals will be taken to upper limit ∞ rather than genome length *L* or chromosome length 1. However, we do consider the chromosomes in considering the total number of ancestral segments, which slightly reduces their mean length.

We first discuss the formation of repeats when *k* is not small (for example *k* ≥ 10). At generation *k* a 1-Morgan chromosome at generation 0 is broken into an expected (1 + *k*) ancestral segments, (Section 2.2). By exchangeability, each has expected length 1/(*k* + 1). For the purposes of this section we will assume that in a large set of *C* chromosomes (e.g. *C* = 3000) there are a large total of *C*(*k* + 1) segments, with independent exponential length distribution ℰ(*k* + 1).

In a segment length *ℓ* the probability of more that 1 crossover is *q*(*ℓ*) = 1 − (1 + *ℓ*) exp(−*ℓ*). Overall the probability an ancestral segment at. depth *k* forms a repeat is

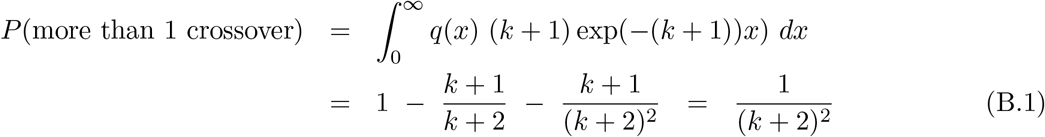

The expected number of new within-chromosome repeat structures formed at generation *k* is

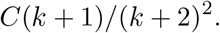

Note we ignore here the simultaneous formation of multiple repeats, as is reasonable for larger *k*.

Segment in which repeats are formed are expected to have lengths *ℓ* longer than average, due to the weighting *q*(*ℓ*). Given a repeat is formed the pdf *f*,(·) of the segment length is

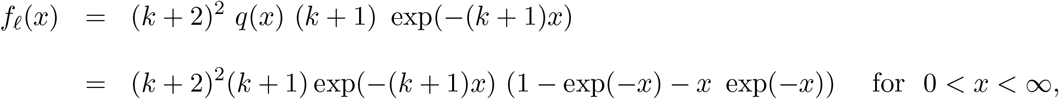

which has mean

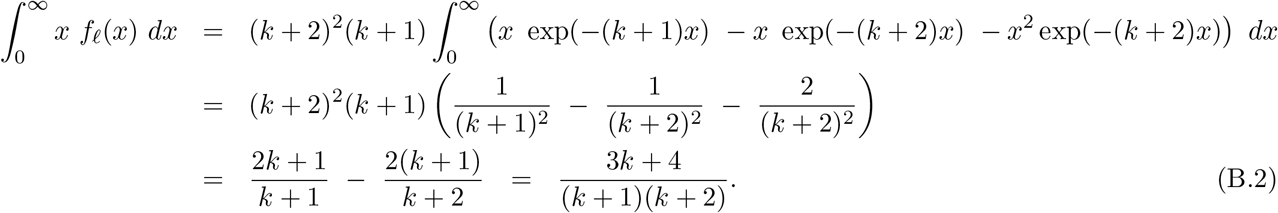

That is, the segments that form into repeats are of the order 3 times longer than a random segment.

The formation of a single repeat by two crossovers in an ancestral segment divides the segment into two (repeated) ancestral segments and an intervening *spacing* segment. Conditional on this 2-crossover event, the positions are a uniform sample on the segment length, so each of the two ancestral segments and the intervening non-ancestral segment have expected length *μ*_*k*_ = (3*k* + 4)/(3(*k* + 1)(*k* + 2)), or very similar to the length of a random ancestral segment: note 1/(*k* + 2) *< μ*_*k*_ *<* 1/(*k* + 1).

**Figure B1:**
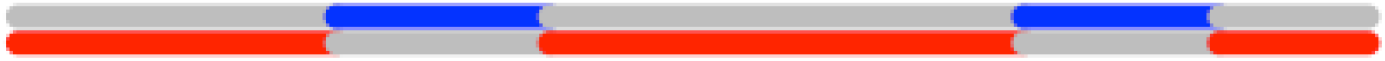
Three repeats formed in two lineages by 4 crossovers in meiosis of an intact chromosome

#### B.2 Formation of multiple repeats in intact chromosomes

Appendix B.1 gives a poor approximation for small *k*. In particular, far too few repeats are formed in the early generations. The main reason is that the above theory ignores the possibility of ancestral segments having three or more crossover events, and hence generating simultaneously 2 or more repeats. Although a full analysis is impractical, we can consider the case of intact chromosomes. That is, all chromosomes at the first generation, and subsequently any chromosome that remains intact.

The probability an intact 1-Morgan chromosome remains intact for *k* generations is

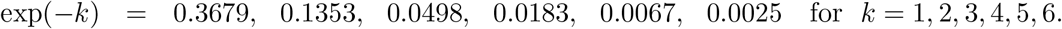

It is then is broken by *m* crossovers with probability

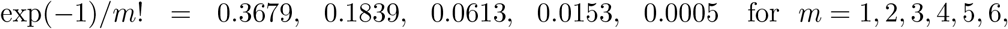

creating (*m* − 1) repeats, for *m* ≥ 2, with mean expected length of intervening non ancestral material (the *spacing*)

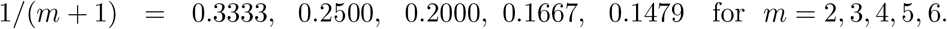

Note that if *m* > 2, these repeats occur in both the two parental chromosomes (Figure B.1). Note also that each of the three spacings corresponds to an ancestral segment in the alternate parental chromosome.

Dividing the total set of 3000 chromosome into those expected to remain intact at each generation, and those that have undergone at least one crossover event, and applying these formulae to the former, gives a far closer approximation to the realizations for the counts of repeats formed in the early generations *k* = 1, 2, 3 (See Figure 15), and a much more accurate estimate of the mean lengths of spacings in repeats formed at each generation (See Figure 16).

#### B.3 Loss of repeats

A repeat is lost when there is a recombination (an odd number of crossovers) in the spacing segment. (Figure 11). Note first that this is a completely independent process from crossovers in ancestral segments at this genomic location: that ancestry has diverged and will be in a different ancestral lineage. Nor does that intervening non-ancestral genome relate to any other chromosome under consideration. Second note that this is a recombination process (odd number of crossovers): an even number of crossovers in any meiosis in the non ancestral spacing will not affect the ancestry of the genome in any way. The probability of recombination is given by the Haldane map function (Equation (1)). There will be no recombination in a length *ℓ* Morgans. with probability

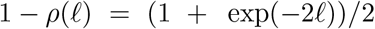

If we assume that the spacing *ℓ* is exponential with mean *μ* a random repeat will survive with probability

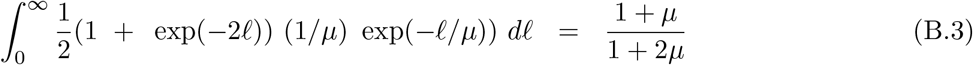

Note that for a given repeat with spacing *ℓ*, the probability of loss remains constant over the generations. However, repeats with larger *ℓ* are lost with higher probability, so that the mean length of the spacing in surviving repeats reduces over the generations, and the probability of survival increases.

If we also approximate the spacing lengths in repeats newly formed by *m* (*m* > 1) crossovers in intact ancestral chromosomes by an exponential, then, from Equation (B.3), each repeat survives the next generation with probability

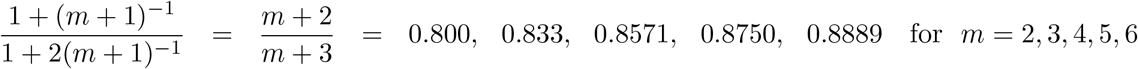

Again, however, the mean spacing lengths will be attenuated with each successive generation. Survival probabilities will increase: loss probabilities will decrease.

If, when a repeat is formed at generation *k* the spacing has pdf *f*_*k*_(*ℓ*) on 0 *< ℓ <* 1 then the spacing pdf for repeats surviving at generation (*k* + *j*) is

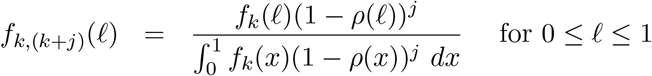

with mean

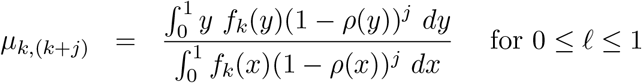

Clearly this distribution is intractable, but when *k* is large, so *ℓ* is small, we can obtain an approximate order-of-magnitude result by assuming exponential distributions: *f*_*k*_(*ℓ*) ∼ ℰ(1*/μ*_*k*_) and

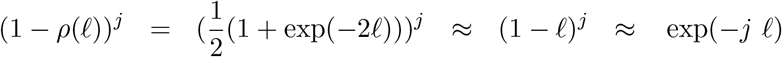

Then

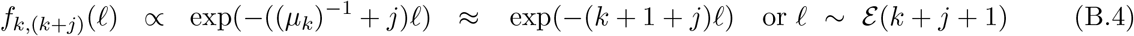

This result, and in particular that *μ*_*k*,(*k*+*j*)_ ≈ 1/(*k* + *j* + 1), explains many of the empirical results in Section 4, in particular those of Figures 16 and 17 and the dependence of expected lengths of surviving spacings primarily on (*k* + *j*) rather than separately on the generation of formation *k*, and survival through the subsequent *j* generations.

